# Biomineral armor in leaf-cutter ants

**DOI:** 10.1101/2020.05.18.102962

**Authors:** Hongjie Li, Chang-Yu Sun, Yihang Fang, Caitlin M. Carlson, Huifang Xu, Ana Ješovnik, Jeffrey Sosa-Calvo, Robert Zarnowski, Hans A. Bechtel, John H. Fournelle, David R. Andes, Ted R. Schultz, Pupa U. P. A. Gilbert, Cameron R. Currie

**Affiliations:** Department of Bacteriology, University of Wisconsin-Madison, Madison, WI 53706, USA; Department of Energy Great Lakes Bioenergy Research Center, Wisconsin Energy Institute, University of Wisconsin-Madison, Madison, WI 53726, USA; Department of Physics, University of Wisconsin-Madison, Madison, WI 53706, USA; Department of Geoscience, University of Wisconsin-Madison, Madison, WI 53706, USA; Department of Entomology, National Museum of Natural History, Smithsonian Institution, Washington, DC 20013-7012, USA; School of Life Sciences, Arizona State University, Tempe, AZ 85287, USA; Department of Medicine, University of Wisconsin-Madison, Madison, WI 53706, USA; Department of Medical Microbiology and Immunology, University of Wisconsin-Madison, Madison, WI 53706, USA; Advanced Light Source Division, Lawrence Berkeley National Laboratory, Berkeley, CA 94720, USA; Departments of Chemistry, Materials Science and Engineering, University of Wisconsin-Madison, Madison, WI 53706, USA

## Abstract

Although calcareous anatomical structures have evolved in diverse animal groups, such structures have been unknown in insects. Here, we report the discovery of high-magnesium calcite [CaMg(CO_3_)_2_] armor overlaying the exoskeletons of major workers of the leaf-cutter ant *Acromyrmex echinatior*. Live-rearing and *in vitro* synthesis experiments indicate that the biomineral layer accumulates rapidly as ant workers mature, that the layer is continuously distributed, covering nearly the entire integument, and that the ant epicuticle catalyzes biomineral nucleation and growth. *In situ* nanoindentation demonstrates that the biomineral layer significantly hardens the exoskeleton. Increased survival of ant workers with biomineralized exoskeletons during aggressive encounters with other ants and reduced infection by entomopathogenic fungi demonstrate the protective role of the biomineral layer. The discovery of biosynthesized high-magnesium calcite in the relatively well-studied leaf-cutting ants suggests that calcareous biominerals enriched in magnesium may be more common in metazoans than previously recognized.

---

Biomineral skeletons first appeared more than 550 million years ago^1–5^, and by the early Cambrian biomineral-based defensive structures had evolved in most extant metazoan phyla, apparently in response to increasing predation pressure^6^. The minerals involved, as well as the biogenic structures they form, are diverse. Calcium-carbonate biomineralization is particularly widespread among metazoans^7^: the hard parts of corals^8^, mollusk shells^9^, stomatopod dactyl club^10^, and sea urchin spines^11^ contain calcium carbonate, as do the light-focusing eye lenses of chitons and brittlestars^12,13^. Magnesium-enriched calcite has been discovered in the central part of the sea urchin tooth, where the increased hardness imparted by magnesium is thought to aid in the grinding of limestone^14–16^. Given the importance of calcareous anatomical structures across metazoan phyla and given that magnesium significantly strengthens such structures, it is surprising that high-magnesium calcite appears to be rare in animals. It is also surprising that, despite the near ubiquity of biogenic mineralization across metazoan phyla and the widespread presence of calcium carbonate in the Crustacea, biomineralized calcium carbonate has so far remained unknown in the most diverse group of animals, the insects, which arose from within the Crustacea^17^. Here we report the discovery of a dense layer of biogenic high-magnesium calcite in the leaf-cutter ants *Acromyrmex echinatior*.

Fungus-growing “attine” ants (tribe Attini, subtribe Attina) engage in an ancient and obligate mutualism with coevolved fungi (order Agaricales), which they cultivate for food. Fungus farming, which has been described as a major transition in evolution^18^, evolved only once in ants around 60 million years ago^18^. Leaf-cutting ants (genera *Acromyrmex* and *Atta*), a phylogenetically derived lineage that arose within the fungus-growing ants around 20 million years ago, harvest fresh vegetation as the substrate on which they grow their fungal mutualists. They are ecologically dominant herbivores in the New World tropics^18,19^ and serve important roles in carbon and nitrogen cycling^20^. A mature leaf-cutter ant colony comprises a “superorganism” with ~100,000 to > 5 million workers, a single queen, and a complex society with a highly refined division of labor based both on worker size and age. In addition to the leaf-cutters, 15 other genera of ants occur within the Attina, all of which grow fungus gardens, form colonies of hundreds to a few thousand workers, and use dead vegetative matter or caterpillar frass rather than fresh leaves and grasses as substrates for their gardens. In addition to the symbiotic association with their fungal cultivars, many fungus-growing ants engage in a second mutualism with Actinobacteria (genus *Pseudonocardia*), which produce antibiotics that help defend the garden from fungal pathogens^21–23^. Fungus-growing ant colonies, containing both fungal crops and immature ant brood, represent a rich nutritional resource for a wide variety of marauding ant species, including army ants and other known “agro-predatory” raiders of ant agriculture. Smaller fungus-growing ant colonies are also subject to attack by the large-sized soldier castes of *Atta* leaf-cutter ants, which use their powerful mandibles to defend their colonies’ territories against other, encroaching ant species^24,25^.

Many species of fungus-growing ants are variably covered with a whitish granular coating, uniformly distributed on their otherwise dark brown cuticles^26^ (Fig. 1a). We report here for the first time that this coating is in fact an outer layer of crystalline mineral covering the ant exoskeleton (Fig. 1b inset) by combining data from *in situ* X-ray diffraction (XRD), electron microscopy, electron backscatter diffraction (EBSD), quantitative electron probe micro-analysis (EPMA), raman and attenuated total reflectance Fourier-transform infrared (ATR-FTIR) spectroscopy. In addition, we conduct synchrotron X-ray PhotoEmission Electron spectro-Microscopy (X-PEEM), *in vitro* synthesis, *in vivo* observation of crystallization and growth through ant-rearing experiment, *in-situ* nanoindentation, ant battle, and infection by entomopathogenic fungi to examine the mechanism of crystal growth and its functional role.

**Fig. 1.**
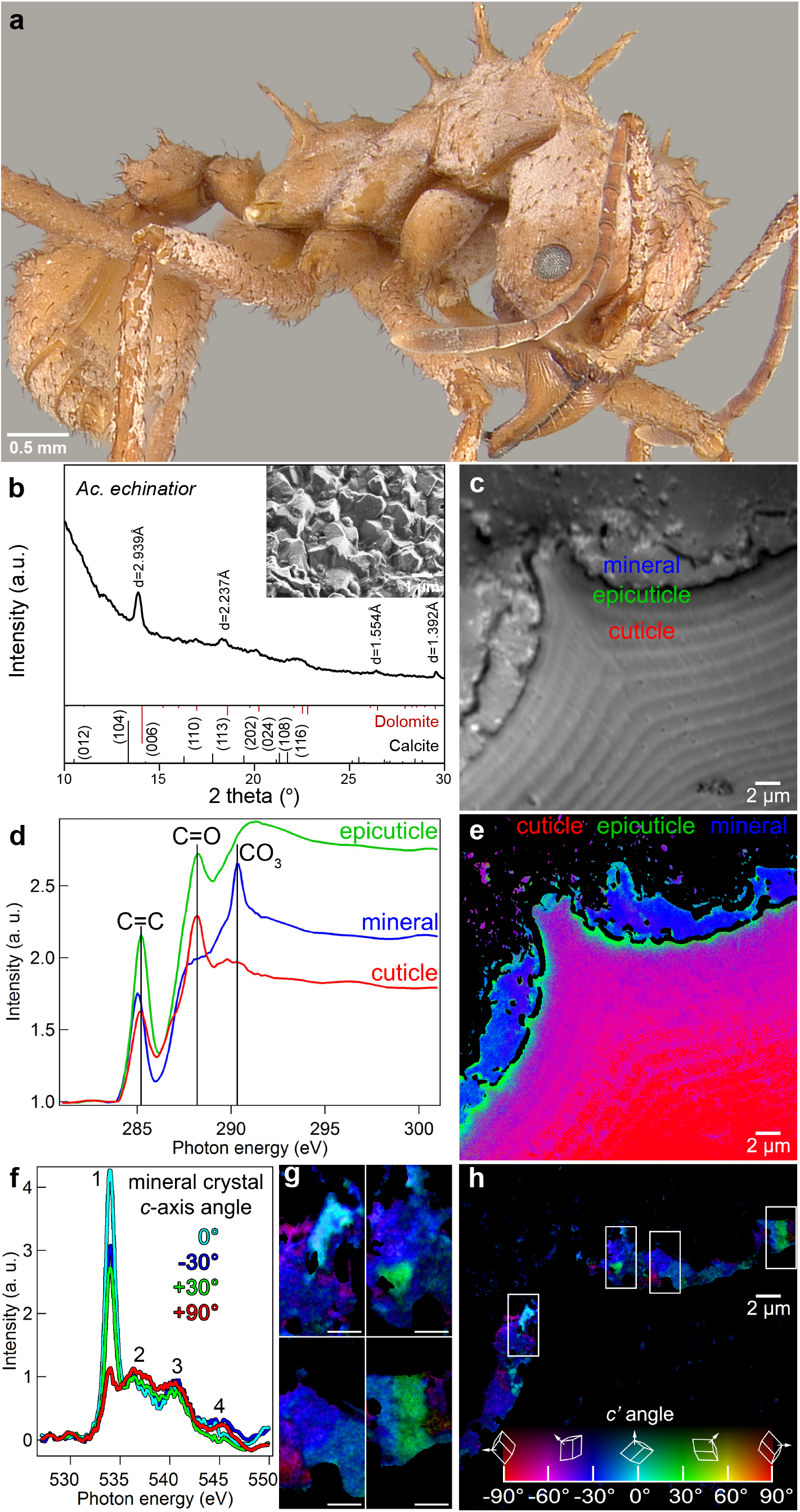
Morphological, structural, and chemical characterization of minerals on the cuticle of the leaf-cutting ant *Acromyrmex echinatior*. **a**, *Ac. echinatior* ant with whitish cuticular coating. **b**, *In situ* XRD analysis identifying the cuticular crystalline layer as high-Mg calcite. Inset: SEM image of ant cuticle with crystalline coating. **c-h**, XANES spectroscopy and mapping with PEEM of cuticular cross-section. **c**, Average of PEEM images acquired across the C K-edge, showing crystalline layer tightly attached to cuticle. Three distinct component spectra were identified in the regions labeled cuticle, epicuticle, and biomineral, from the most internal part of the ant (bottom right of micrograph) to the outer surface. **d**, Normalized component spectra extracted from the corresponding labeled regions. Characteristic peaks are marked, including the 285.2 eV (C=C), 288.2 eV (C=O) and 290.3 eV (carbonate) peaks. **e**, Component map where each pixel is colored according to the chemical components it contains. Black pixels are masked areas containing epoxy or gaps. Faint carbonate components within the cuticle and epicuticle were emphasized by enhancing the blue channel 5×, thus this is a semi-quantitative map. A fully quantitative RGB component map is presented in Supplementary Fig. 23. Individual maps of each component are presented in Supplementary Fig. 24, clearly showing an increasing gradient of carbonates towards the surface in the cuticle. **f**, O K-edge spectra extracted from the mineral crystals correspondingly colored in the Polarization-dependent Imaging Contrast (PIC) maps in g and h. **g**, Magnified PIC maps for the regions represented by boxes in the complete PIC map in h. **h**, PIC map quantitatively displaying the orientations of the mineral crystals’ *c*-axes in colors. This map was acquired from the same area shown in c and e at precisely the same magnification. These are interspersed high- and low-Mg calcite, and heterogenous at the nanoscale. Biomineral crystals do not show preferred orientations, but are randomly oriented. High-magnesium calcite in carbon spectra is identified by the carbonate peak at 290.3 eV, which occurs in all carbonates, amorphous or crystalline. The O spectra in d clearly indicate crystallinity, and their lineshape indicates a mixture of high-magnesium calcite or Mg-bearing calcite.

## Results and discussion

### Morphological, structural, and chemical characteristics of epicuticular minerals

Microscopic imaging of polished cuticular cross-sections of the leaf-cutting ant *Acromyrmex echinatior* reveals a clear interface between this crystalline layer and the ant cuticle (Fig. 1c). This layer is brighter than the cuticle in backscattered electron (BSE) mode scanning electron microscopy (SEM) (Supplementary Fig. 1 and 2), indicating that it consists of heavier elements and that it is continuously distributed, covering nearly the entire integumental surface. Energy-dispersive X-ray spectroscopy (EDS) characterization of the cuticular coating further indicates that the crystalline layer contains significant amounts of magnesium and calcium (Supplementary Fig. 3 a-f), suggestive of a Mg-bearing calcite biomineral. X-ray diffraction (XRD) analysis confirms the high-magnesium calcite composition of the biomineral layer in *Ac. echinatior,* as indicated by the *d*-spacing of (104) peak at 2.939 Å (Fig. 1b and Supplementary Table 1). Quantitative electron probe micro-analysis (EPMA) reveals a magnesium concentration of 32.9±2.7 mol % (Supplementary Table 2). Using bright-field transmission electron microscopy (TEM), selected area electron diffraction (SAED), TEM-EDS, Raman and Attenuated Total Reflectance Fourier Transform Infrared (ATR-FTIR) spectra, we further confirm the biomineral is a high-magnesium calcite with chemically heterogeneous crystals and with no observable Ca-Mg ordering (Supplementary Fig. 4 and 5). Extensive XRD analyses of *Ac. echinatior,* including both lab-reared and field-collected workers from Panama and Brazil, confirms the consistent presence of high-magnesium calcite in quantities of 23–35 mol% MgCO_3_ (Supplementary Table 2 and 3).

The mineral-cuticle interface of *Acromyrmex echinatior* was investigated using synchrotron X-ray PhotoEmission Electron spectro-Microscopy (X-PEEM) at the Advanced Light Source (Lawrence Berkeley National Laboratory, Berkeley, CA) (Fig. 1c)^8^. Distinct X-ray Absorption Near Edge Structure (XANES) spectra occur at the carbon K-edge for each of three regions: cuticle, epicuticle, and mineral layer (Fig. 1d; mapped as spectral components in Fig 1e). The C spectra for the mineral layer show a strong carbonate peak at 290.3 eV (Fig. 1d). The oxygen K-edge spectra extracted from the ant mineral layer indicate that the carbonate crystals are crystalline with a strong crystal orientation dependence of peak 1 at 534 eV (Fig. 1f).

Polarization-dependent imaging contrast (PIC) mapping^27,28^ across the mineral layer, in which color quantitatively displays the orientation of the crystal *c*-axes, indicates that crystals are randomly oriented (Fig. 1, g and h). The width of peak 2 in all O spectra (Fig. 1f) indicates a mixture of phases with high- and low-Mg concentrations^29^. This chemical heterogeneity is consistent with the XRD data (Fig. 1b), with electron microprobe analyses (Supplementary Table 2) backscatter diffraction (EBSD) results (Supplementary Fig. 6), and with the magnified PIC map regions (Fig. 1g).

Unlike the typical chitin spectrum of insect epicuticle, the *Acromyrmex echinatior* XANES epicuticular spectrum is consistent with a protein-enriched insect epicuticle^30^. Protein hydrolysis of the cuticular layers verifies that the epicuticle is proteinaceous (Supplementary Fig. 7). Further, the epicuticular spectrum shows a very intense peak at 285.2 eV (Fig. 1d), which, based on its energy position and its symmetric line shape, suggests that the epicuticle contains one or more phenylalanine (Phe) enriched proteins^31^. High Performance Liquid Chromatography (HPLC) amino acid profiling of the protein layer of the *Ac. echinatior* epicuticle confirms the presence of phenylalanine in the ant cuticle (Supplementary Table 4).

### *In vitro* high-mg calcite synthesis

To assess whether epicuticular proteins mediate the precipitation of high-magnesium calcite in *Acromyrmex echinatior*, we performed synthetic biomineralization experiments in which the cuticle of *Ac*. *echinatior* was incubated in saturated carbonate solutions with a [Mg^2+^]/[Ca^2+^] ratio of 5 at ambient conditions^32^ (Fig. 2a and Supplementary Fig. 8). In these *in vitro* experiments, nanocrystal aggregates precipitated on the epicuticle of *Ac*. *echinatior* (Fig. 2, b and c), and were identified as anhydrous high-magnesium calcite by XRD and EDS analyses (Fig. 2d and Supplementary Fig. 9). As a negative control, we performed the same *in vitro* mineralization experiments using the cuticle of the leaf-cutter ant *Atta cephalotes,* which belongs to the sister genus of *Acromyrmex,* does not have a biomineral cuticular layer, and has different cuticular structures (Supplementary Fig. 10). We found that only aragonite crystals formed, mainly on the hairs of *At*. *cephalotes* and almost never on the epicuticle (Supplementary Figs 11 and 12), indicating direct precipitation from solution since aragonite is the favorable crystalline precipitate in high Mg conditions^33^. In control experiments using *Ac*. *echinatior* epicuticles either treated with KOH to hydrolyze surface proteins or coated with a 10 nm platinum layer to disable protein function, only aragonite crystals formed (Fig. 2d, Supplementary Fig. 9). Interestingly, in synthetic biomineralization experiments using cuticle from different developmental stages (pupae to fully mature adult workers), we found that only mature worker epicuticles catalyze the precipitation of high-magnesium calcite (Fig. 2e), consistent with the presence of a more substantial protein layer in mature workers indicated by SEM examination (Supplementary Figs. 13 and 14). These *in vitro* synthesis results suggest that the protein layer in the epicuticle of *Ac*. *echinatior* and the unusual morphological structures on the cuticles of the ants catalyze the low-temperature nucleation and growth of magnesium-rich calcite on the epicuticles of mature workers of *Ac. echinatior*.

**Fig. 2.**
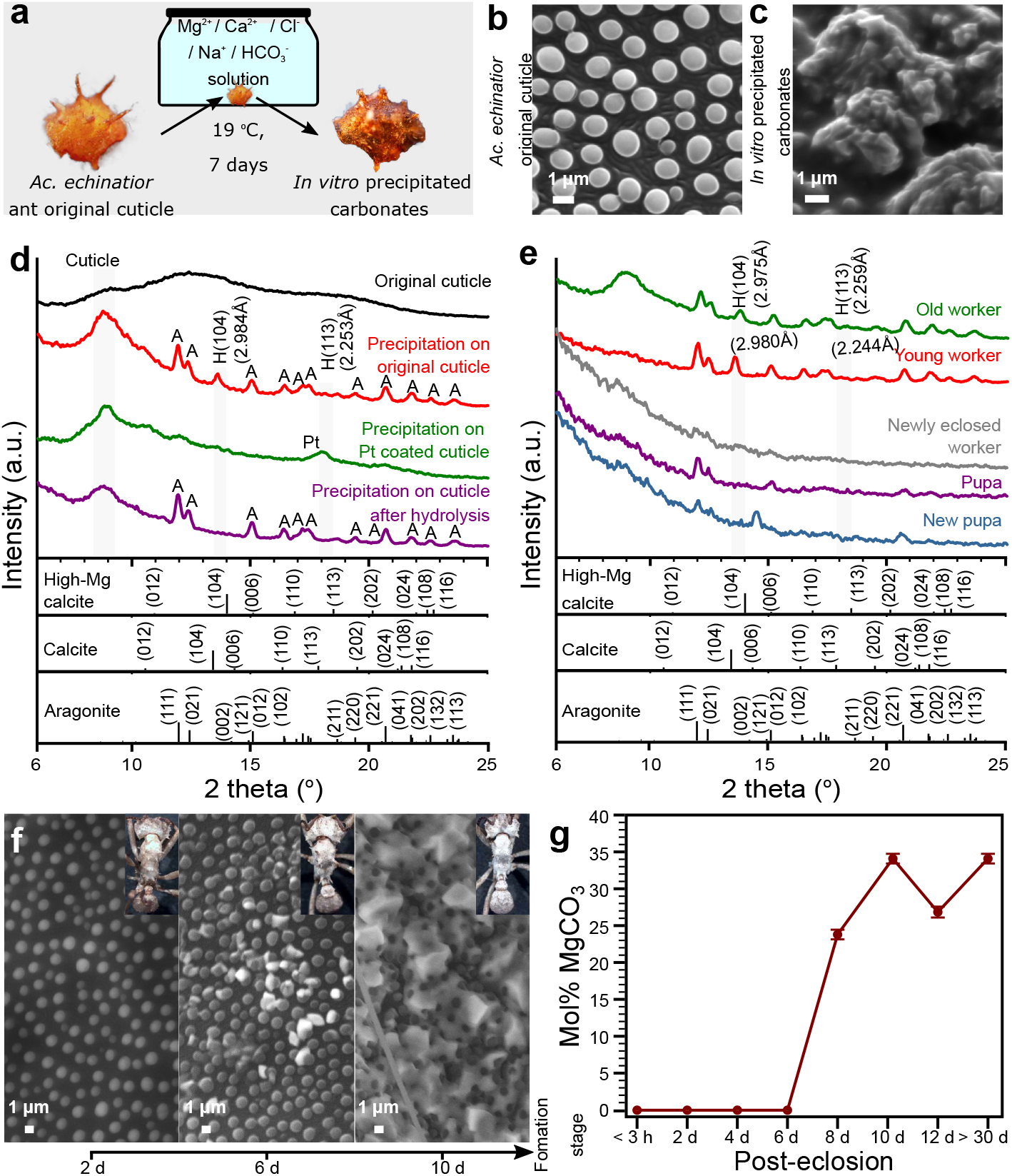
Mineral precipitation on the cuticle of the leaf-cutting ant *Acromyrmex echinatior* in both *in vitro* cuticle synthetic studies and ant-rearing experiments. **a**, Scheme of *in vitro* mineralization experiment using *Acromyrmex echinatior* leaf-cutting ant cuticles as templates for biomineralization. **b** and **c**, Pre- and post-incubation SEM images showing the original, uncoated cuticle (b) and the cuticle covered by a layer of precipitated carbonate (c) after incubation in Mg^2+^/Ca^2+^/Cl^−^ /Na^+^/HCO_3_^−^ solution for 7 days at 19 ^o^C. **d**, XRD patterns of, from top to bottom, an uncoated ant cuticle, a cuticle after incubation in Mg^2+^/Ca^2+^/Cl^−^/Na^+^/HCO_3_^−^ solution, a platinum-coated cuticle incubated in Mg^2+^/Ca^2+^/Cl^−^/Na^+^/HCO_3_^−^ solution, and a cuticle after KOH protein hydrolysis incubated in Mg^2+^/Ca^2+^/Cl^−^/Na^+^/HCO_3_^−^ solution. H: high-magnesium-calcite, A: aragonite, Pt: platinum. **e**, XRD patterns of cuticles of ants representing different developmental stages, ranging from (from bottom to top), a newly formed pupa to an older worker, after incubation in Mg^2+^/Ca^2+^/Cl^−^/Na^+^/HCO_3_^−^ solution. **f**, Environmental scanning electron micrographs (eSEM) of ant epicuticles taken over a 10-day time series, from immediately after eclosion from pupa to adult (left), to 10 days post-eclosion (right), showing the formation of the biomineral layer over time. **g**, Estimated magnesium concentration of the biomineral layer during 30 days of ant development based on the XRD *d*(104) value according to Graf and Goldsmith (1956)^39^, showing the rapid integration of magnesium from days 6 to 8 and the continued presence of high magnesium content for up to 30 days.

### *In vivo* crystallization and growth of high-mg calcite

To explore the developmental timing of biomineral formation on the epicuticles of *Acromyrmex echinatior* workers, we conducted rearing experiments. Twenty pupae at the same developmental stage were collected, randomly sorted into two groups of ten, and reared to callow adults, then one worker from each of the two groups was collected every second day and analyzed by XRD and eSEM (Fig. 2, f and g). No biomineral layer was visible nor detected with XRD on workers 0 to 6 days after eclosion from the pupal to adult stage. In contrast, 8 days after eclosion visible and XRD-detectable high-magnesium calcite was present on workers. Magnesium was rapidly integrated into the calcareous biomineral in these older workers, with XRD measurements of mol% MgCO_3_ reaching ~35% within 2 days after the initiation of biomineralization on individual worker ants (i.e., from days 6 to 8 after eclosion; Fig. 2g).

Three independent lines of evidence indicate that epicuticular biomineral crystals are ant-generated rather than adventitiously precipitated from the environment or generated by bacteria. First, in both C component maps (e.g. Fig. 1e) and PIC maps (e.g. Fig. 1g) the magnesium-rich calcite crystals outside the epicuticle are space-filling, a characteristic of biominerals formed by eukaryotes^34^. Second, magnesium-rich calcite biominerals are spatially co-localized with epicuticular protein(s), which are likely involved in biomineral formation, consistent with the absence of biomineral formation in *in vitro* synthesis experiments in which ant epicuticles were either coated with platinum or excluded. Third, the ant rearing experiments were carried out in sterile, clean Petri dishes, eliminating the possibility of biominerals acquired from external sources.

### Mechanical protection of epicuticular high-mg calcite

It is plausible that epicuticular high-magnesium calcite enhances the structural robustness of the ant exoskeleton, providing better defense for ants engaged in ‘wars’ with other ants or under attack from predators or parasites. To test this hypothesis, we first quantified the increase in hardness conferred by the protective biomineral layer using *in-situ* nanoindentation in an SEM (Fig. 3a, Supplementary Fig. 15 and Supplementary Video 1). Since the surface of the exoskeleton is not flat, conventional nanoindentation could not be used, whereas *in-situ* nanoindentation with real-time microscopic imaging allowed near-perpendicular contact of the probe tip with the surface (Supplementary Fig. 15). Typical non-biomineralized ant cuticle, made primarily of chitin, has a hardness of H ~ 0.73±0.04 GPa (Fig. 3a and Supplementary Fig. 16). In contrast, when high-magnesium calcite and cuticular layers are combined, the composite structure has a greater than two-fold increase in hardness (1.55±0.48 GPa, compared to cuticle alone of 0.73±0.04 GPa) (Fig. 3a and Supplementary Fig. 16 and 17). Given that the biomineral layer has an average thickness of 2.3 μm and that it overlays a cuticle with an average thickness of 33.5 μm, this more than two-fold increase in hardness is conferred by only a 7% increase in cuticle thickness (Supplementary Fig. 18). Additional *in-situ* nano-mechanical testing of the cuticles of *Atta cephalotes* ants, which do not have a biomineral layer, as well as of other common insects, including a beetle (*Xylotrechus colonus*) and a honey bee (*Apis mellifera*), produced similar hardness values in the range of 0.4–0.7 GPa (Fig. 3a and Supplementary Fig. 16) as they are all mainly made of chitin. The nano-mechanical measurements indicate that the biomineralized layer substantially hardens the exoskeleton of *Ac. echinatior*, consistent with the hypothesis that the biomineral layer functions as protective armor.

**Fig. 3.**
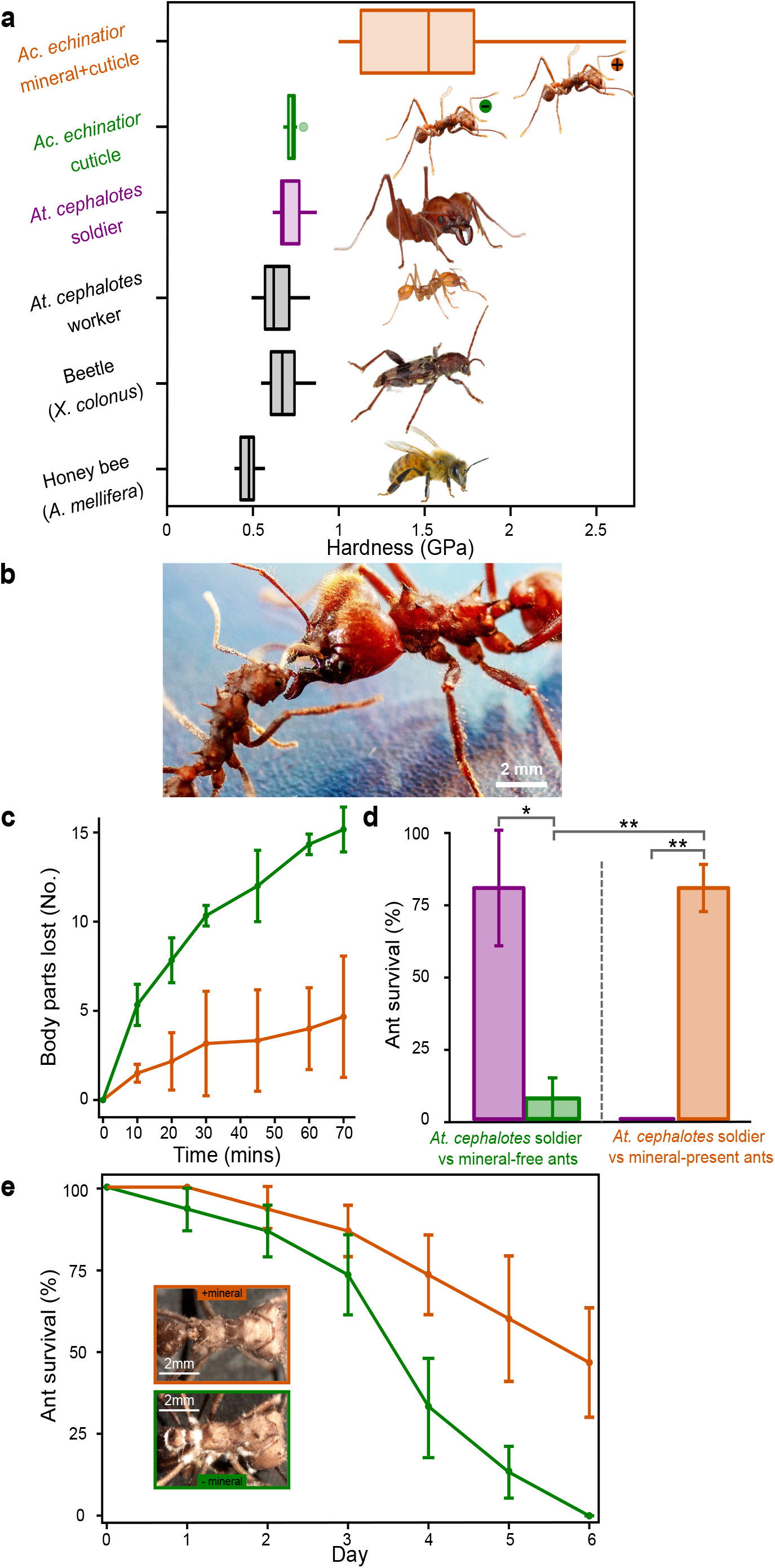
Mechanical protection afforded by the epicuticular mineral layer. **a**, Quantitative nano-mechanical properties of insect cuticles, including honey bee (*Apis mellifera*), beetle (*Xylotrechus colonus)*, leaf-cutting ants (*Atta cephalotes* worker, *Atta cephalotes* soldier, and *Acromyrmex echinatior* worker without biomineral) and *Ac. echinatior* ant worker with biomineral epicuticular layer, measured by an *in-situ* nanoindenter with a cube-corner probe. **b-d**, Aggressive interaction between three *Ac. echinatior* workers (with/without biomineral, respectively) and *Atta cephalotes* soldier. **b**, *Ac. echinatior* worker (left) aggressively interacts with *Atta cephalotes* soldier (right). **c**, In aggressive encounters with *Atta cephalotes* soldiers, *Ac. echinatior* workers with biomineral armor (orange) lose substantially fewer body parts (i.e., legs, antennae, abdomen, and head) compared to *Ac. echinatior* worker without biomineral (green). **d**, Survivorship of *Ac. echinatior* workers without (green) and with (orange) biomineral armor in aggressive encounters with *Atta cephalotes* soldiers (purple). Asterisks indicate significant differences via two-sample *t* test (\**P* < 0.05; \*\**P* < 0.001). **e**, Survivorship curves of *Ac. echinatior* worker with and without an epicuticular biomineral layer exposed to the entomopathogenic fungus *Metarhizium*. The inset images show more substantial fungal growth and emergence from biomineral-free workers.

To further test the role of the biomineral as protective armor, we exposed *Acromyrmex echinatior* major workers with and without biomineral armor to *Atta cephalotes* soldiers in ant aggression experiments designed to mimic territorial ‘ant wars’ that are a relatively common occurrence in nature^24,35,36^. In direct combat with the substantially larger and stronger *At. cephalotes* soldier workers (average body length of 10.4 mm and a head capsule width of 6.1 mm, compared to major *Ac. echinatior* body length of 6.4 mm and head capsule width of 2.9 mm) (Fig. 3b), ants with biomineralized cuticles lost significantly fewer body parts (Fig. 3c and Supplementary Fig. 19) and had significantly higher survival rates compared to biomineral-free ants (Fig. 3d, Supplementary Video 2 and 3). Further, in direct aggression experiments in which biomineral-armored *Ac. echinatior* workers were pitted against *At. cephalotes* soldiers, all of the *At. cephalotes* soldiers died, whereas only a few such deaths occurred when *Atta* soldiers were pitted against biomineral-free ants. SEM examination of biomineral-armored *Ac. echinatior* ants after combat with *Atta cephalotes* soldiers showed significantly less damage to their exoskeletons (Supplementary Fig. 20). Notably, biomineral armor is present in mature major workers, which forage outside of the nest, further indicating that epicuticular high-magnesium calcite is critical in a highly competitive environment (Supplementary Figs 21 and 22). These results, taken together, are consistent with a role for epicuticular high-magnesium calcite as armor that defends workers from aggressive interactions with other ants.

Biomineral armor could also help protect ants from pathogens. In a series of experiments, we focused on entomopathogenic fungi, which establish infection by penetrating the insect exoskeleton and have significant impacts on survival. We exposed propleural plate of *Ac. echinatior* major worker ants with and without biomineralized exoskeletons to the spores of the entomopathogenic fungus *Metarhizium anisopliae* (Ascomycota, Hypocreales). Compared to biomineral-free workers, major workers with biomineralized exoskeletons were significantly more resistant to infection. Specifically, we found that a majority of ants without biominerals died from infection within 4 days (1.0 ± 0.4 and 0 ± 0 ants survived to 4 and 6 days, respectively), whereas an average of 2.2 ± 0.4 and 1.4 ± 0.5 (out of 3 individuals per sub-colony over 5 sub-colonies) ants with biominerals survived to 4 and 6 days, respectively (Fig. 3e). All ants succumbed to infection within 6 days. Examination of workers without biominerals exposed to *M. anisopliae* revealed substantial fungal growth and emergence (Fig. 3e inset).

The biota of the Ediacaran period (635 to 541 million years ago) included organisms of known and unknown phylogenetic affinities that lived in oceans with a high ratio of magnesium to calcium. Most were soft-bodied, but some possessed rudimentary skeletons composed either of aragonite (a form of calcium carbonate) or, notably, of high-magnesium calcite^6^. Around 550 million years ago, coinciding with a shift in the Earth’s oceans to significantly lower magnesium-to-calcium ratios, metazoans with strongly calcified internal and external skeletons appeared, including most familiar modern phyla. In spite of its strengthening properties, the enrichment of calcareous structures with high concentrations of magnesium in Cambrian and modern metazoans has until now remained only known from a very small plate within the tooth of sea urchins. The ability of fungus-growing ants to facilitate the formation of magnesium-rich biominerals on their epicuticles is thus surprising. Further, given that fungus-growing ants are among the most extensively studied tropical insects, our finding raises the intriguing possibility that high-magnesium calcite, and perhaps even partially ordered dolomite, biomineralization may be more widespread than previously suspected.

Fungus farming in ants originated ~60 million years ago in South America when a hunter-gatherer ancestor irreversibly committed to subsistence-scale cultivation of fungal crops for food^37^. The transition to industrial-scale agriculture occurred ~20 million years ago with the origin of the ecologically dominant leaf-cutting ants, in which colony populations are orders of magnitude greater in size and in which physically distinct worker castes enable complex division of labor, paralleling the similar importance of agriculture in driving the expansion of human populations and the elaboration of human social systems^38^. Further paralleling human agriculture, the fungal cultivars of the ants are highly susceptible to pathogens and the ants have responded, in part, by evolving associations with antibiotic-producing bacteria to protect their crops^22^. Early sedentary human agricultural settlements represented rich resources that were highly susceptible to marauding bands of human raiders, leading to the development of multiple modes of defense, including specialized warrior castes, fortified cities, weapons, and protective armor^38^. Here we show that, in another striking parallel with agriculture-driven human cultural evolution, fungus-growing ants have evolved biomineralized armor that serves, at least in part, to protect them from other ants, including other fungus-growing ants in disputes over territory and “agropredatory” ants that are known to raid their colonies and to consume their gardens and brood.

## Methods

### Photoemission electron microscopy (PEEM)

*Acromyrmex echinatior* ants were freeze-dried prior to PEEM sample preparation. The heads of the ants were then detached and embedded in Epofix epoxy (EMS, Hatfield, PA), ground with SiC sandpapers, polished with Al_2_O_3_ suspensions of 300 nm (MicroPolish II, Buehler, Lake Bluff, IL) and 50 nm (Masterprep, Buehler, Lake Bluff, IL) particle sizes^8,40^. 22 g/L Na_2_CO_3_ saturated solution was added regularly onto the pad during grinding and polishing to prevent carbonate dissolution, and the Al_2_O_3_ suspensions were also dialyzed against 22 g/L Na_2_CO_3_ saturated solution^41^. The samples were re-embedded to fill as much as possible the interior of the ants and the gap between mineral and epoxy, and then the polishing procedures were repeated. After final polishing, the samples were rinsed with ethanol and gently wiped with TexWipe Cotton (Texwipe, Kernersville, NC), air dried, and coated with 1 nm Pt on the areas to be analyzed and 40 nm Pt around it^42^.

For C K-edge spectra, PEEM stacks were acquired by scanning across 280-320 eV range with 0.1-eV step between 284 and 292 eV, and 0.5-eV step elsewhere, resulting in 145 images per stack^43^. For O K-edge spectra, PEEM stacks were acquired by scanning across 525-555 eV range with 0.1-eV step between 530 and 545 eV, and 0.5-eV step elsewhere, resulting in 181 images per stack^44,45^. The images were stacked and processed with GG Macros in Igor Pro 6.37^46^.

For PIC mapping, a stack of 19 images were acquired by fixing the photon energy at the O K-edge π* peak (534 eV) and changing the X-ray polarization from horizontal to vertical with a 5° step^40,47,48^. Colored PIC maps were then produced using Igor Pro 6.37 with GG Macros^46^.

### Masking the component map

The component map in Fig. 1e was masked using an image of the same region acquired in SEM in backscattered electron (BSE) mode. Unfortunately, in both the PEEM average image in Fig. 1c and in the BSE image, the gray levels in the embedding epoxy and those in the cuticle are similar. Therefore, there is no rigorous and quantitative method to select one but not the other. We used Adobe Photoshop and the Magic Wand tool with a tolerance of 30 to select all of the cuticle and deleted all those pixels from a black mask. The brighter mineral and all of the mineral debris deposited on the epoxy were then selected using the Magic Wand and a tolerance of 50 on the BSE image. These were also deleted from the same black mask. The black pixels in the BSE image correspond to gaps between the cuticle and the epoxy, or holes between mineral crystals, those black pixels were remained black in the black mask. The bright mineral debris is presumably an artifact of polishing, as they appear both in PEEM and SEM images and are spectroscopically identified without a doubt as mineral. These were also removed from the mask and therefore displayed in Fig. 1e, as removing them would have been an artifact. The BSE image was warped to correspond correctly to the PEEM image using Adobe Photoshop and specifically the Puppet Warp tool.

### Obtaining component spectra

We extracted single-pixel spectra from the cuticle, the epicuticle, and the mineral regions. These were identified as the only 3 reliable components that were spectroscopically distinct from one another and not linear combinations of other components. The single-pixel spectra from the same material were extracted from each stack, aligned in energy, and averaged. The averaged spectrum was then normalized to the beamline I_0_ curve, acquired with precisely the same energy steps.

The 3 spectra were then aligned between 280.0 and 283.7 eV. The cuticle and epicuticle spectra were shifted in energy so that the first peak was at 285.2 eV for chitin and proteins, following Cody et al. 2011^30^, whereas the mineral spectra was shifted in energy so that the last peak, characteristic of carbonates, was at 290.3 eV, following Madix and Stöhr^49,50^. The cuticle spectrum is identical to that published by Cody et al. 2011 obtained from scorpion cuticle, and interpreted as chitin. The spectrum has a peak at 285.2 corresponding to C=C in aromatic carbon, a shoulder at ~287 eV, and a peak at 288.2 eV corresponding to C=O in chitin. The epicuticle spectrum shows the characteristics C=C of aromatic amino acids (tyrosine, tryptophan, and phenylalanine)^31^, a shoulder at 287.6 eV corresponding to C-H aliphatic carbon, and a sharp peak at 288.2 eV corresponding to carboxyl group (C=O) in the peptide bonds of all proteins. Compared to the spectra in tyrosine and tryptophan, the phenylalanine spectrum has a more symmetric peak at 285.2 eV, allowing us to assign this peak to phenylalanine in the epicuticle^31^. The C=O occurs at the expected 288.2 eV (Myers et al. 2018)^43^. These normalized and averaged spectra were then adopted as “component spectra”, displayed in Fig. 1d, and used to obtain a component map in Fig. 1e.

### Component mapping

The extracted, averaged, normalized, and aligned component spectra were made references by multiplying the I_0_. Spectrum in each pixel of the stack was analyzed and best-fitted to a linear combination of the component references: cuticle, epicuticle, and mineral. The resulting component proportion maps were exported as a gray level image and combined by the Merge Channel function in Adobe Photoshop, which became a fully quantitative RGB image (Supplementary Fig. 22). Individual component distribution maps were presented in Supplementary Fig. 23. For Fig. 1e, we enhanced the blue channel by adjusting the midtone value in Levels 5 times greater than the other two channels, to emphasize the presence of mineral in the cuticle and epicuticle.

### *In vitro* synthesis

All synthesis experiments were carried out in sealed plastic bottles at 19°C for 7 d. Solutions were prepared by dissolving 50 mM MgCl_2_·6H_2_O, 10 mM CaCl_2_·2H_2_O, and 50 mM NaHCO_3_ with pH buffer to ~8.0 with NaOH to simulate modern seawater chemistry. Solution were mixed for 20 minutes and then divided into 100 mL bottles with ant exoskeleton (Supplementary Fig. 5). Peptide synthesis experiments with 1 mM, 5 mM, and 10 mM of phenylalanine peptide (H-Phe-Phe-Phe-OH) (Bachem, CA) were mixed into solutions without ants. All vessels during the experiments have been washed with deionized water and pretreated with 6M hydrochloric acid to prevent carbonate contamination. Filtered solution and ants were airdry for XRD and SEM characterization.

### Scanning electron microscopy (SEM) and electron backscatter diffraction (EBSD)

Scanning electron microscopy (SEM) were done using a Hitachi S3400 at 15 kV. Images were obtained in both variable pressure and vacuum mode. Energy-dispersive x-ray spectroscopy (EDS) and electron backscatter diffraction (EBSD) were carried out using an AZtecOne system with silicon-drift detector from Oxford instruments. Samples were coated with 5 nm Pt coating. Phases used in EBSD are constructed based on Mg-poor (a = 4.990Å, c = 17.062Å; Graf, 1961), Mg-medium (a = 4.920Å, c = 16.656Å; calculated), Mg-rich (a = 4.850Å, c = 16.250Å) regions. However, given the EBSD is not particularly sensitive to unit-cell parameter differences, chemical heterogeneity from EBSD is only qualitative.

### Transmission electron microscopy (TEM)

TEM measurements were carried out using a Philips CM200UT TEM instrument operating at 200 kV acceleration voltage with 0.5 mm spherical aberration (Cs) and a point resolution of 0.19 nm. Images and electron diffraction were collected with a CCD camera and analyzed with Gatan DigitalMicrograph software. Samples that have been previously examined by XRD and SEM were rinsed with ethanol and DI water to remove glue residue. Samples were then crushed in an agar mortar, suspend in acetone, and drop onto Lacey/ carbon 200 mesh copper grid. Composition of phases were confirmed with energy-dispersive X-ray spectroscopy (EDS) and analyzed with Thermo Noran software.

### *In situ* X-ray diffraction (XRD) analyzes

*In situ* X-ray diffraction (XRD) were performed using a Rigaku Rapid II X-ray diffraction system with Mo Kα radiation. This XRD instrument use a 2-D image-plate detector for signal collection and integrated using Rigaku’s 2DP software. XRD were run at 50 kV and a 100-μm diameter collimator. Whole fresh ant samples were glued onto American Durafilm Kapton® tube with vacuum grease. Ant samples were then spin around phi and oscillate on omega. Synthesized powder samples were sealed in Kapton tube and run with fixed omega and phi spin. Refinements for phase percentage and unit-cell parameter were run using Jade 9.0 software with American Mineralogist Crystal Structure Database (AMCSD) and the PDF-4+ database from the International Centre for Diffraction Data (ICDD). Disordered dolomite reference was constructed based on unit cell parameter of a disordered dolomite with 50 mol.% MgCO3 and powder diffraction pattern calculated by CrystalMaker built-in CrystalDiffract software.

### Sectioning and transmission electron microscopy (TEM)

Ants for sectioning and transmission electron microscopy were fixed in cold 2% glutaraldehyde in Na-cacodylate buffer. Postfixation was done in 2% osmium tetroxide and specimens were subsequently dehydrated in a graded acetone series. Specimens were embedded in Araldite and sectioned with a Reichert Ultracut E microtome. Semithin 1-μm sections for light microscopy were stained with methylene blue and thionin. Double-stained 70-nm thin sections were examined in a Zeiss EM900 electron microscope.

### Quantitative electron probe micro-analysis (EPMA)

The carbonate EPMA data were acquired with a CAMECA SXFive FE electron probe in the Cameca Electron Probe Lab in the Department of Geoscience at the University of Wisconsin-Madison. Operating conditions were 7 kv and 10 nA (Faraday cup), using a focused beam. A low accelerating voltage was used to shrink the analytical volume to less than 300 nm. Peak counting time was 10 seconds, with background acquired for 10 seconds. Mg Ka was acquired with a TAP crystal and Ca Ka with an LPET crystal. The standard used as Delight Dolomite. Automation and data reduction utilized Probe for EPMA (Probe Software), Carbon and oxygen were accounted for in a robust procedure in the Probe for EPMA software: oxygen was calculated based upon stoichiometry to the measured Mg and Ca, with carbon calculated relative to that oxygen value (1:3), with this being iterated several times within the Armstrong/Love Scott matrix correction. The resulting values were then evaluated for actual accuracy, based upon two criteria: a non-normalized analytical total close to 100 wt% (~98-102 wt%), and for a formula basis of 3 oxygens, the carbon formula value being close to 1.00 (~.99-1.01). With these conditions met, the determined compositions were deemed acceptable.

### Raman and attenuated total reflectance Fourier-transform infrared (ATR-FTIR) spectroscopy

Ant mineral samples for Raman experiments were prepared by bleaching the freeze-dried ant samples in 8.25% NaClO commercial bleach for 24h at room temperature to remove the exposed organic materials (*7*). Raman spectra were collected using a LabRam Raman microprobe (JY Horiba, Inc.) equipped with a Microscope (Olympus DX41, 50X and 100X objectives) and a 633 nm laser. Spectra were acquired with a CCD camera behind a spectrometer (the accumulations and integration time varied). The ant carbonate powders were dropped on a microscope slide just before individual measurement.

For ATR-FTIR, freeze-dried ant samples were used directly. ATR-FTIR data collection was conducted on a Perkin-Elmer 1720x spectrometer according to manufacturer’s instructions.

### Rearing experiments

In total of twenty worker pupae of *Ac. echinatior* at the same developmental stage and its ~10g fungus-garden were collected and randomly sorted into two groups of ten and maintain them in chambers (diameter 6 cm, height 4 cm) with wet cotton. Reared to callow workers (around 3 hours) and followed by one worker from each of two groups was collected every second day. The fresh ant samples were subjected to XRD analyses immediately and followed by Environmental scanning electron microscope examination (eSEM). For eSEM, an FEI QUANTA 200 eSEM (FEI Company) was used. Ants were placed directly onto the eSEM stub and examined without any preparation (i.e., samples were not fixed or coated for this analysis). All samples were analyzed at 5.0 torr, 3.0 spot size, and 4 °C.

### Ant sample preparation for SEM analyses of cuticular structure

Worker ant cuticles from *Ac. echinatior* at different developmental stages (pupae to fully mature adult workers) and *At. cephalotes* were immediately fixed with 4% (vol/vol) formaldehyde and 1% glutaraldehyde at 22°C RT overnight. Samples were then washed with PBS and treated with 1% osmium tetroxide for 30 min at 22°C. Samples were subsequently washed with a series of increasing ethanol dilutions (30 to 100% [vol/vol]), followed by critical point drying and coating with 1-nm platinum. Scanning electron microscopy (SEM) of samples was performed using a LEO 1530 microscope to investigate the cuticular structure.

### High Performance Liquid Chromatography (HPLC) analysis

Dissected ant cuticle samples were placed in 100 μl 25% TFA containing 10 mM DTT and were hydrolyzed at 110°C for 24 h. Hydrolyzed samples were then dried at 45°C under stream of nitrogen and resuspended in 50 mM HCl. Amino acids were then converted into respective fluorescent derivatives using *o*-phthalaldehyde (OPA) (Agilent #5061-3335). Briefly, 5 μl sample aliquot was added to 20 μl of 40 mM potassium tetraborate buffer (pH 9.8) followed by addition of 5 μl OPA, mixed gently and another 40 μl water was added. The mixture was filtered through a 0.45 μm cellulose acetate 4-mm syringe filter (Nalgene #171-0045). Freshly prepared samples were immediately subjected to HPLC analysis.

Amino acids were analyzed using a modified method described elsewhere^51^. The apparatus used was a custom-built dual analytical/semi-preparative Shimadzu system consisting of a SIL-20AC autosampler, a CBM-20A system controller, two LC-20AR pumps, a C50-20AC oven, a PDA S10-M20A detector, and a CPP-10Avp detector. Chromatographic separation of OPA-derivatized amino acids was performed using an Agilent ZORBAX Eclipse AAA column (4.6mm x 150 mm x 3.5 μm; Agilent #963400-902) coupled with a ZORBAX Eclipse AAA Analytical Guard Column (4.6 mm x 12.5 mm x 5 μm; Agilent #820950-931) heated at 40°C. The gradient elution was applied using 40 mM sodium phosphate dibasic buffer (pH 7.8) as solvent A and a mixture of acetonitrile, methanol, and water (45:45:10, v/v/v) as solvent B. HPLC-grade acetonitrile and methanol were supplied from Fisher Scientific and were used without further purification. The optimum separation of amino acids was obtained using the following gradient program: 0% B for 1.9 min, then increase to 57% B up to 28.10 min followed by increase to 100% B up to 38.60 min, then hold at 100% B till 47.30 min, and decrease to 22.3% B up to 48.20 min and down to 0%B till 60 min. The flow rate was 1 mL/min. Aliquots of 10 μL standards/samples were injected at 0.5 min and amino acids were detected at the maximum wavelength of 338 nm with a 4 nm bandwidth. Retention times as well as spectral information given by the PDA detector were used for peak identification. Calibration curves of individual and mixed amino acids were prepared using either 250 pmol stocks of corresponding individual amino acids or a 250 pmol amino acid standard mix (Agilent #5061-3331). Quantification was performed using calibration curves of the respective amino acid standards.

### Biomineral-free *Ac. echinatior* ant generation

Biomineral-free *Ac. echinatior* ants were generated using a sub-colony setup. Sub-colonies were set up in small (diameter 6 cm, height 4 cm) clear plastic containers. After sterilizing containers for at least 20 minutes using UV light, cotton moistened with distilled water was placed at the bottom to help provide humidity. A small (width 4.12 cm, length 4.12 cm and height 0.79 cm) weigh boat (Fisher catalogue #08-732-112) was placed on top of the wet cotton, and then 0.1 g of fungus garden, 2 minor workers, and a major worker pupa being reared to derive a biomineral-free adult (n = 10 sub-colonies). A ∼1 cm2 leaf fragment of pin oak (*Quercus palustris*) was added 24 hours or more after pupa eclosion for the ants to cut and incorporate into the fungus garden. We monitored sub-colonies daily to record eclosion date for the major worker pupa until 14–21 days after eclosion. Then we performed environmental Scanning Electron Microscopy (eSEM) and XRD on a subset of the ants to confirm mineral absence. Meanwhile, we established that *Ac. echinatior* ants could grow mineral normally in this sub-colony with the addition of 2 major workers (n = 10 sub-colonies) and other colony components were maintained as above.

### Nanomechanical testing

Nanoindentation tests were carried out using a Bruker Hysitron PI-85 SEM Picoindenter in a Zeiss Leo 1550VP SEM at the Wisconsin Centers for Nanoscale Technology, UW-Madison. The samples were tested using a cube-corner probe with a basic quasistatic trapezoid load-controlled function, where the maximum load was 500 μN, the hold time was 2 sec, and the loading/unloading rate was 100 μN/sec^52^. SEM imaging was done in high vacuum using accelerating voltage of 3 kV with secondary electron mode. In order to simulate the defense mechanism of the actual ant exoskeleton, we tested the combination of ant mineral and ant cuticle by indenting from the outside in, as illustrated in Supplementary Fig. 13. The samples for testing the combination of ant mineral and ant cuticle were prepared by slicing through the transverse plane of the head of the ant to allow probing only on the flatter top part of the head. All samples were then attached firmly to carbon tape on an SEM stub. The “combination” sample was additionally pressed carefully with tweezers to ensure good attachment and flatness. The indentation data and corresponding SEM video were analyzed to ensure: 1) that the correct contact point of the probe with the surface was chosen, 2) there was no movement of the sample during indentation, and 3) the load-displacement curve was smooth and there was no abrupt pop-out or discontinuity. Based on these criteria, 13, 13, 10, and 12 valid data points were selected for *Ac*. *echinatior* ant mineral plus cuticle and *Ac*. *echinatior* ant cuticle cross-section, respectively. Measurements on other insect cuticles were also done by indenting from the outside in following the same analyzing criteria, with which 15, 13, 15, and 12 valid data points were selected for Atta soldier ant cuticle, Atta worker ant cuticle, beetle elytra, and honeybee cuticle, respectively. All the valid points were then used to obtain the hardness values^53^ presented in Fig. 3a. Additional hardness comparison of the mineral phases alone were done on polished cross-sections of geologic dolomite and *Ac*. *echinatior* mineral. Cross-section samples for were prepared using the same embedding and polishing procedures as described for the PEEM samples. Representative load-depth curves in Supplementary Fig. 14 were selected from the data point closest to the averaged hardness value in each sample.

### Aggressive experiments between *At. cephalotes* soldier and *Ac.echinatior* workers

We confronted one major *At. cephalotes* soldier and three mineral-present/mineral-free workers of *Ac*. *echinatior*^54^. The experiment was replicated 5 times under reduced light in 9cm Petri dishes. Survival of ants were counted after 3 hours of confrontation.

Using time lapse setting of an iPad Pro 2018, we recorded aggressive encounters between *At. cephalotes* soldier and *Ac*. *echinatior* workers (Video S2 and S3). For the aggressive experiments with mineral-present ants, the video was started when *At. cephalotes* soldier was placed in the petri dish with the *Ac*. *echinatior* ants. Video was stopped when the soldier ant was killed and the *Ac*. *echinatior* worker was able to separate from the solider ant. The video is a total of 42 minutes filmed in time lapse at 120 times its speed reducing the video to 21 seconds in length, followed by editing it down to 10% of its speed using Premiere Pro resulting in a 3:27 minutes video. For the aggressive experiments with mineral-free ants, video was started when the soldier ant was placed into the petri dish, video was stopped after all 3 *Ac*. *echinatior* ants were killed by the solider ant. The video was recorded for 1 hour in time lapse at 120 times its speed reducing the video to 30 seconds in length, followed by editing down to 10% of its original speed resulting in a video that is 5:00 minutes.

### Entomopathogenic fungi infection

Using *Ac. echinatior* major worker ants present and absent mineral, we conducted infection experiments using the entomopathogenic fungus *Metarhizium anisopliae* (Ascomycota, Hypocreales)^55^. In brief, every three ants were placed into an individual Petri dish with a ring of moist cotton. Then, each ant’s propleural plate was inoculated with 0.5 μl *Metarhizium* spores of ca. 1.00 × 10^7^ conidiospores ml^−1^ suspension + 0.01% Tween 20 by using a micropipette under dissecting microscope. The experiment was replicated 5 times and inoculated with control solution of sterile, deionized water + 0.01% Tween 20^56^. The survival of ants was monitored every 24 h post-treatment for 6 days.

## Acknowledgement

We thank U. P. Agarwal and S. A. Ralph from US Forest Products Laboratory for the Raman spectroscopy measurements; B. Schneider and R. Noll for expert help conducting SEM work; J. Morasch for assistance with nanoindentation measurements; E. Okonski for ant imaging and laboratory assistance; and R. J. Massey for assistance with microtome.

## Author contributions

Study design: H.L., C.-Y.S., P.U.P.A.G., C.R.C. Experimental design and supervision: H.L., C.-Y.S., T.R.S., P.U.P.A.G., C.R.C. Data collection and analysis: H.L., C.-Y.S., Y.F., C.M.C., H.X., A.J., J.S-C., R.Z., H.A.B., J.H.F., D.R.A., T.R.S., P.U.P.A.G., C.R.C. Initial draft: H.L., C.R.C., T.R.S. C.-Y.S., P.U.P.A.G. Final version: All authors.

## Data and materials availability

All data is available in the main text or the supplementary materials.

## Supplementary Information

Supplementary Text

Supplementary Tables 1-4

Supplementary Figures 1-26

Supplementary Videos 1-3

References (57-59)

