## Supplemental Materials for "Biomineral armor in leaf-cutter ants"

**This PDF file includes:**

Supplementary Text
Supplementary Tables 1 to 4
Supplementary Figs. 1 to 26
Captions for Videos 1 to 3
Supplementary References

**Other Supplementary Materials for this manuscript include the following:**

Supplementary Videos 1 to 3

### SUPPLEMENTARY TEXT

**Caste and age differences in presence of mineral armor.** Comparisons of the worker castes (minor and major workers) in *Ac. echinatior* revealed that major workers have mineral, whereas in the minor workers the mineral is absent (Supplementary Figs 21 and 22). These differences are correlated with the caste differences in task, which major workers have more activity for foraging outside while minor workers mostly hang over around the fungus-garden. We further compare white pupae, brown pupae, newly molted worker, middle-age worker and old workers. The mineral is only present in middle-age worker and old workers by XRD analyses. These differences across castes and age-groups in the presence/absence of mineral, together with only old major worker forage outside nest, can be expected to correlate with mechanical protection.

**Age differences in epicuticular dynamic changes and capacity of mineral precipitation.** Comparisons of the major workers across developmental stages (pupae, brown pupae, newly molted worker, middle-age worker and old workers) revealed that only mature workers (middle-age workers and old workers) enable to precipitate the high-magnesium-calcite (Fig. 2e). Meanwhile, our SEM studies show distinct difference in epicuticular surface, matrix accumulate gradually along with development (Supplementary Fig. 14). These differences are well consistent to the epicuticle development, which strong support the phenylalanine-rich protein layer mediated Mg-bearing calcite precipitation.

**Sources of Magnesium and Calcium ion.** SEM image of *Ac. echinatior* revealed abundant secretions scatter on the epicuticle (Supplementary Fig. 25, a and b), and the after wash show the specialized tubercle structures (Supplementary Fig. 25 c), which our previous studies show is connected to an internal glandular cell[19,21,57]. Therefore, these are glandular secretions. Our further EDS analysis show the glandular secretions enriched with Magnesium and Calcium ions (Supplementary Fig. 25, d and e).

***In vitro* synthesis using Phe peptides.** To assess whether the phenylalanine catalyze precipitation of Mg-bearing calcite, we performed the synthetic biomineralization experiments using the phenylalanine-rich peptides at 1 mM, 5 mM, and 10 mM concentration. The filters are further subjected to XRD and we found that aragonite crystals were exclusively formed (Supplementary Fig. 26). Given peptides chain length and structure showed variant efficiencies in biomineralization studies[58,59], our results suggested that potential structural peptide for Mg-bearing calcite formation.

**Supplementary Table 1. Biogenic and geologic sampling list represented in this work.**

| Sample name | Collection Year | Collection Country | Deposited |
|---|---|---|---|
| Geologic Calcite | 2018 | Brazil | Physics Dept, UW-Madison |
| Sea urchin (*Strongylocentrotus purpuratus*) | 2018 | USA | Physics Dept, UW-Madison |
| Attine ant (*Acromyrmex echinatior*) | 2019 | Panama | Bacteriology Dept, UW-Madison |
| Attine ant (*Atta cephalotes*) | 2018 | Panama | Bacteriology Dept, UW-Madison |
| Beetle (*Xylotrechus colonus*) | 2019 | USA | Entomology Dept, UW-Madison |
| Honey bee (*Apis mellifera*) | 2018 | USA | Bacteriology Dept, UW-Madison |
| Geologic Dolomite | 1921 | USA | Geology Dept, UW-Madison |

**Supplementary Table 2. Unit cell parameters and quantitative electron probe micro-analysis (EPMA) of biogenic and geologic samples.**

| | Unit cell parameters (Å) | | | | mol% $MgCO_3$ | |
|---|---|---|---|---|---|---|
| | $d_{(104)}$ | $d_{(015)}$ | a | c | XRD estimation* | EPMA |
| Calcite | 3.032 | - | 4.990 | 17.062 | 0 | - |
| Sea urchin spine (*S. purpuratus*) | 3.019 | - | 4.962 | 16.973 | 5.8 | 3.2±0.7 |
| *Ac. echinatior* | 2.939 | - | 4.859 | 16.431 | 33.3 | 32.9±2.7 |
| | 2.935 | - | 4.846 | 16.430 | 34.7 | 30.9±1.3 |
| | 2.939 | - | 4.852 | 16.444 | 33.3 | 30.9±1.9 |
| Dolomite | 2.889 | 2.532 | 4.811 | 16.110 | 50.4 | 46.6±0.6 |

*Note the magnesium contents in crystalline calcite based on the XRD $d_{(104)}$ value according to Graf and Goldsmith (1956)[39].

**Supplementary Table 3. XRD analyses of *Acromyrmex echinatior* worker ants from Panam and Brazil.** Note the magnesium contents in crystalline calcite based on the XRD $d_{(104)}$ value according to Graf and Goldsmith (1956)[39].

| Ant species | Collection Country | $d_{(104)}$ | %mol MgCO3 |
|---|---|---|---|
| *Ac. echinatior* | Panama | 2.947 | 30.2 |
| *Ac. echinatior* | Panama | 2.960 | 25.8 |
| *Ac. echinatior* | Panama | 2.968 | 23.0 |
| *Ac. echinatior* | Brazil | 2.960 | 26.0 |
| *Ac. echinatior* | Brazil | 2.956 | 27.4 |

**Supplementary Table 4. HPLC-determined percentage amino acid profiles of the *Ac. echinatior* cuticle.** Different developmental stages are included for comparison.

| Sample | ASP | GLU | SER | HIS | THR | ARG | ALA | TYR | GLY | CYS | VAL | MET | PHE | ILE | LEU | LYS |
|---|---|---|---|---|---|---|---|---|---|---|---|---|---|---|---|---|
| Fresh pupa | 8.1 | 13.1 | 6.6 | 1.9 | 3.2 | 5.3 | 12.5 | 6.9 | 18.9 | 1.4 | 4.5 | 2.3 | 3.1 | 2.5 | 9.3 | 0.3 |
| Pupa | 8.4 | 8.4 | 6.5 | 1.0 | 4.1 | 2.5 | 16.4. | 3.5 | 8.6 | 19.1 | 5.3 | 0.4 | 2.3 | 3.3 | 7.8 | 2.3 |
| Callow worker | 6.9 | 7.3 | 6.8 | 0.5 | 4.6 | 3.3 | 21.3 | 3.1 | 17.5 | 7.3 | 5.7 | 0.3 | 2.2 | 3.8 | 8.5 | 1.0 |
| Young worker | 2.7 | 3.8 | 8.6 | 1.7 | 3.0 | 3.8 | 20.5 | 2.2 | 33.8 | 4.5 | 3.9 | 0.1 | 1.7 | 2.5 | 6.8 | 0.3 |
| Old worker | 2.4 | 3.9 | 4.9 | 7.6 | 2.3 | 3.7 | 19.6 | 2.9 | 35.9 | 0.7 | 3.8 | 0.3 | 2.5 | 2.6 | 7.0 | 0.0 |

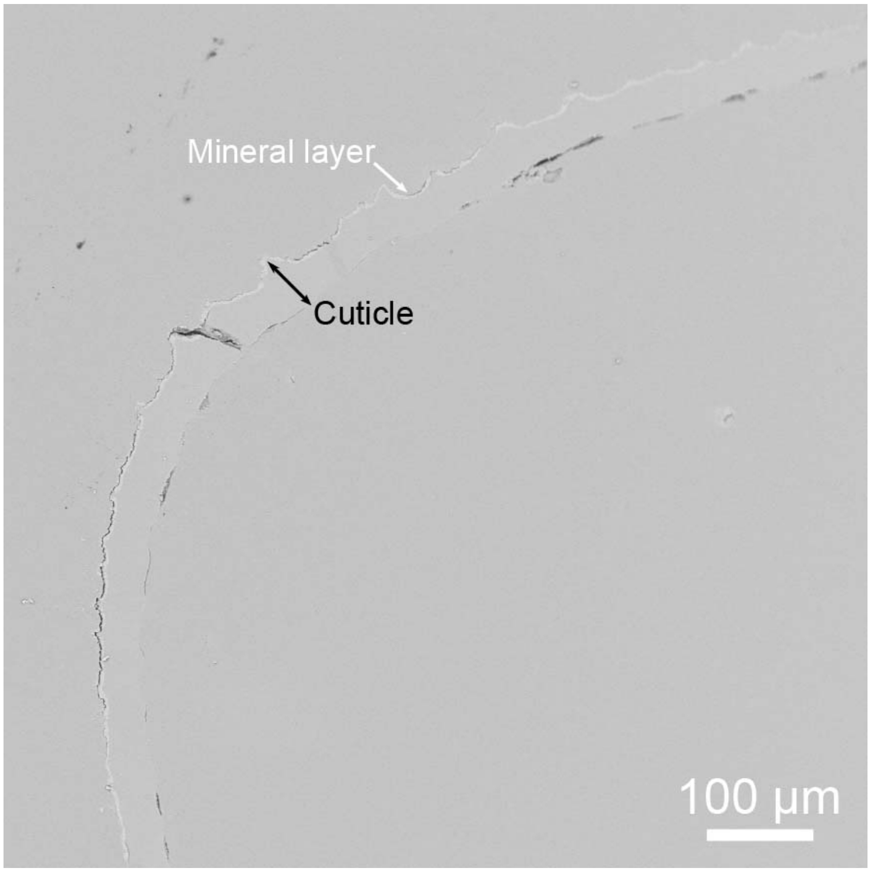


**Fig. S1. Backscattered electron (BSE) image of a polished cuticular cross-sections of the leaf-cutting ant *A. echinatior.*** Polished cross-section of the cuticle reveals the continuity and relative thickness of the mineral layer to the entire cuticle. This layer is brighter than the cuticle in backscattered electron (BSE) mode scanning electron microscopy (SEM), indicating that it consists of heavier elements and is continuous and covering nearly the entire surface.

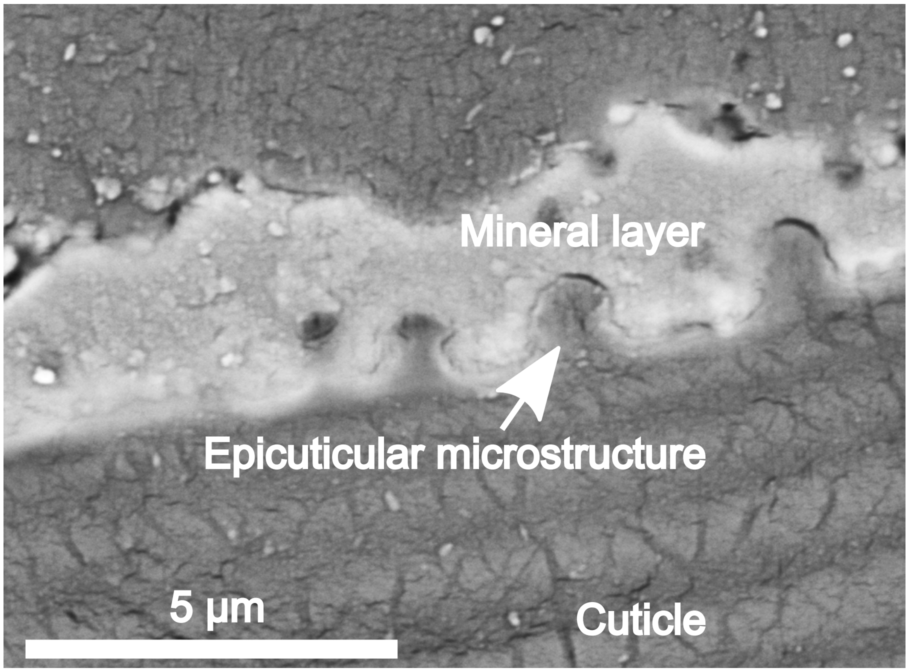


**Supplementary Fig. 2. Backscattered electron (BSE) image of a polished cuticular cross-sections of the leaf-cutting ant *Ac. echinatior*.** Polished cross-section of the cuticle reveals the continuity and relative thickness of the mineral layer to the entire cuticle. This layer is brighter than the cuticle in backscattered electron (BSE) mode scanning electron microscopy (SEM), indicating that it consists of heavier elements.

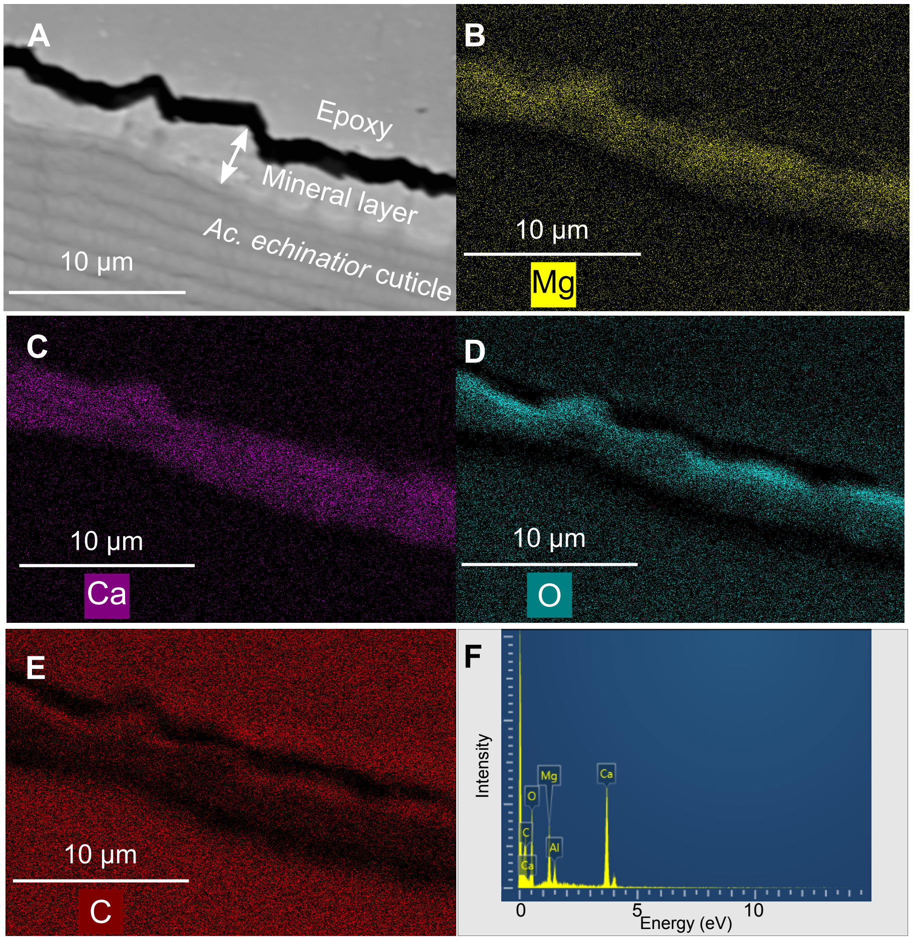


**Supplementary Fig. 3. Energy-dispersive X-ray spectroscopy (EDS) analyses of** a **polished cuticular cross-section of the leaf-cutting ant *Ac. echinatior*. (A)** BSE SEM image of the polished cross-section, showing the mineral-cuticle interface. EDS mapping revealed much higher **(B)** Mg and **(C)** Ca concentrations in the mineral layer**. (D)** O and **(E)** C are present in both the cuticle and the mineral layer. **(F)** EDS spectra shows the average chemical compositions of the ant mineral crystals. Note that the black band in (A) is a gap due to embedding and polishing, and the aluminum in the EDS in (F) resulted from the $Al_2O_3$ particles used for polishing.

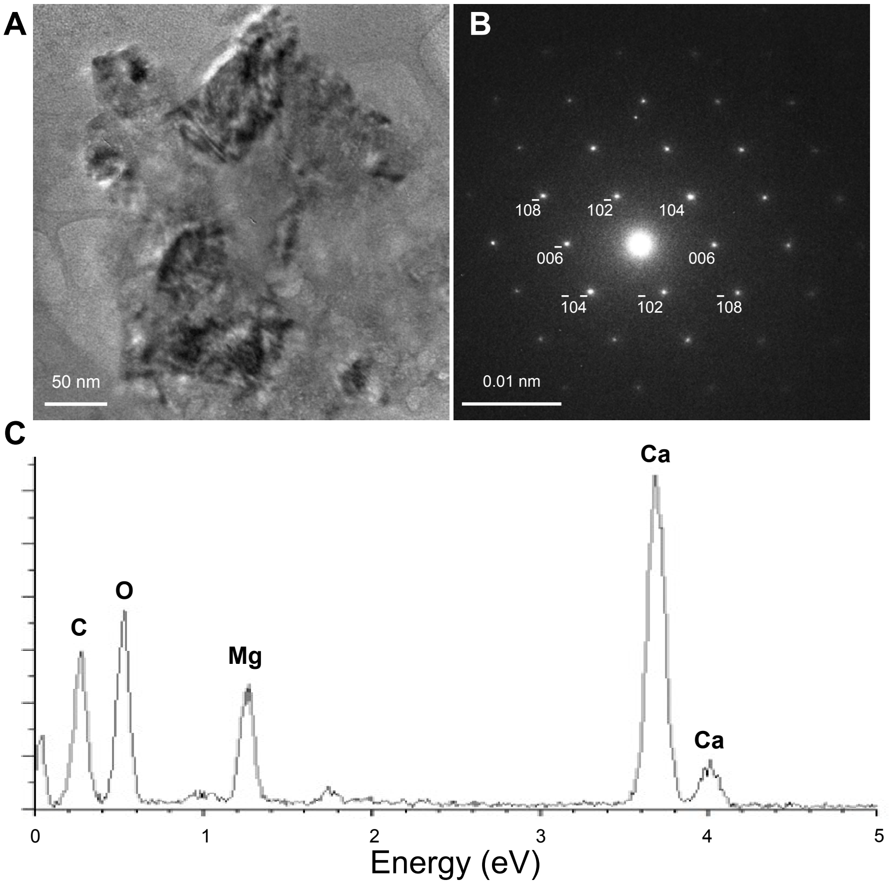


**Supplementary Fig. 4. Morphological and surface features of *Ac. echinatior* ant mineral crystals.** Bright field TEM micrographs, corresponding selected area electron diffraction (SAED) patterns and energy-dispersive X-ray spectroscopy **(**EDS) of *Ac. echinatior* ant mineral crystals. **(A)** TEM shows that crystals are euhedral and less than 100 nm in sizes. Strong strain contrast indicates individual crystals are heterogeneous and composed of many nanodomains, in which each nanodomain has slightly different composition and displays small angle boundaries. **(B)** SAED pattern doesn't show Ca-Mg ordering. **(C)** The EDS shows the average chemical compositions of the ant mineral crystals on the TEM.

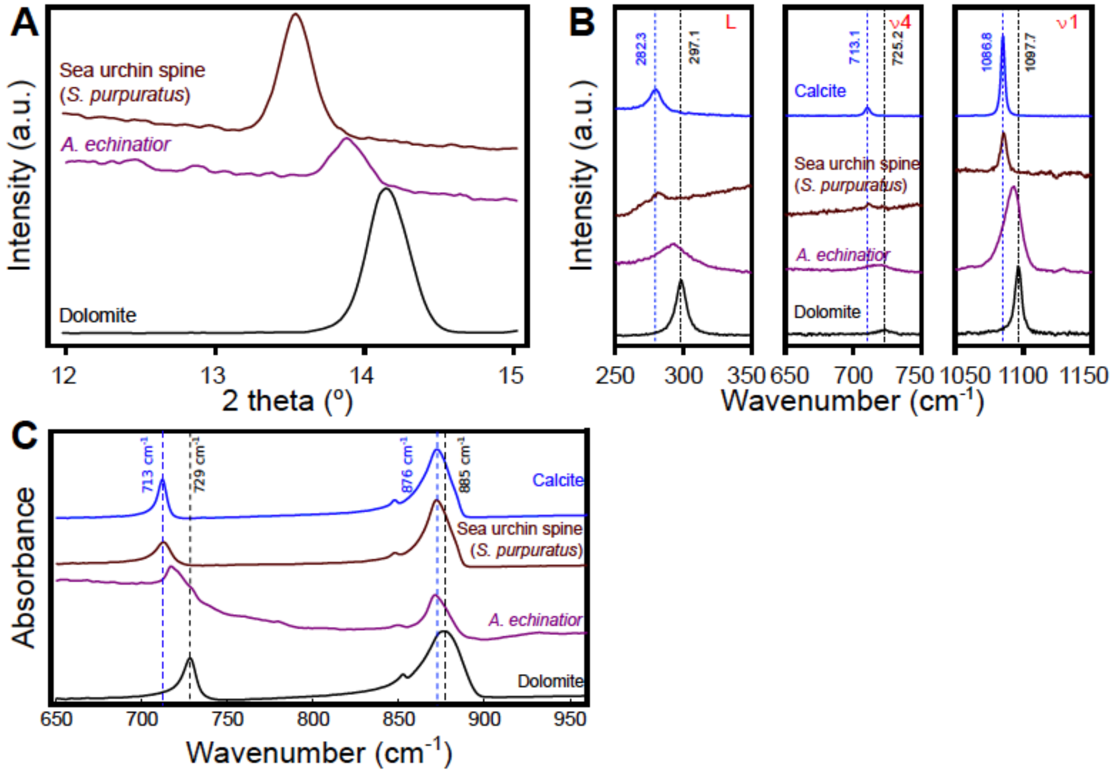


**Supplementary Fig. 5. XRD, Raman and FTIR spectra of biogenic carbonates from sea urchin spine, *A. echinatior* ants, as well as geogenic calcite and dolomite.**

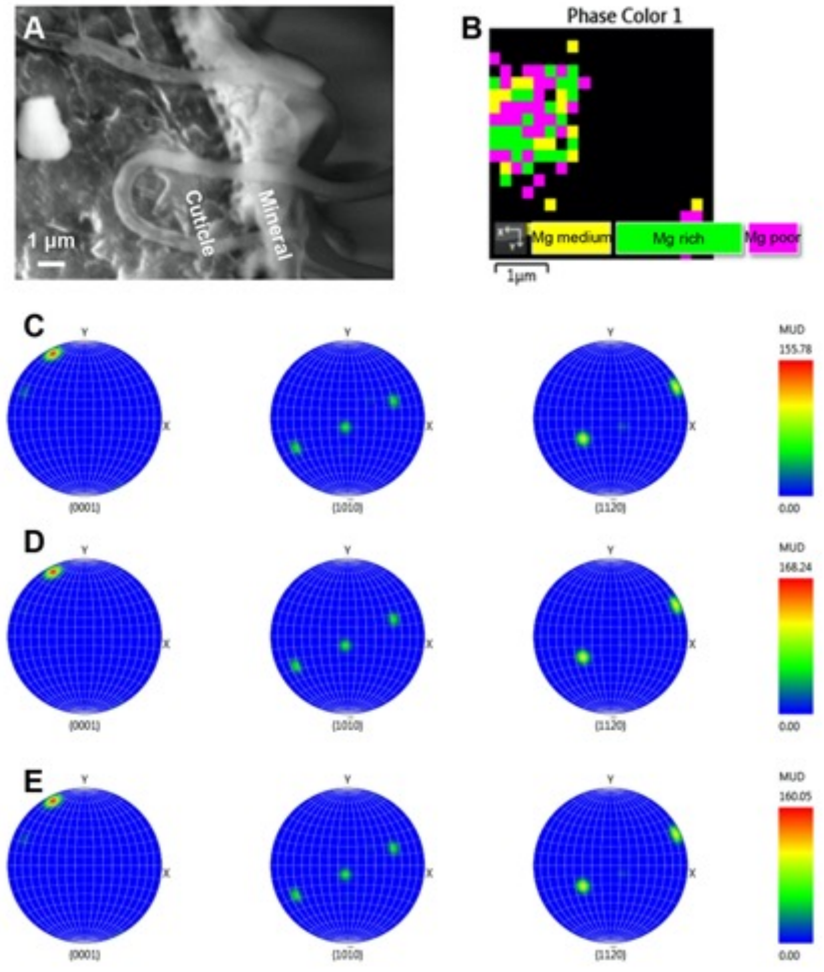


**Supplementary Fig. 6. The chemical heterogeneity of ant mineral crystals. (A)** A **backscattered electron (**BSE) SEM image showing mineral crystals on exoskeleton of head of *Ac. echinatior* major worker. The image clearly shows mineral crystals on the exoskeleton. **(B)** The electron backscatter diffraction (EBSD) result showing heterogeneous MgCO3 content in the mineral crystal form (A). **(C, D** and **E)** Pole figures of the Mg medium region, Mg rich region and Mg poor region in the crystal with same crystallographic orientation.

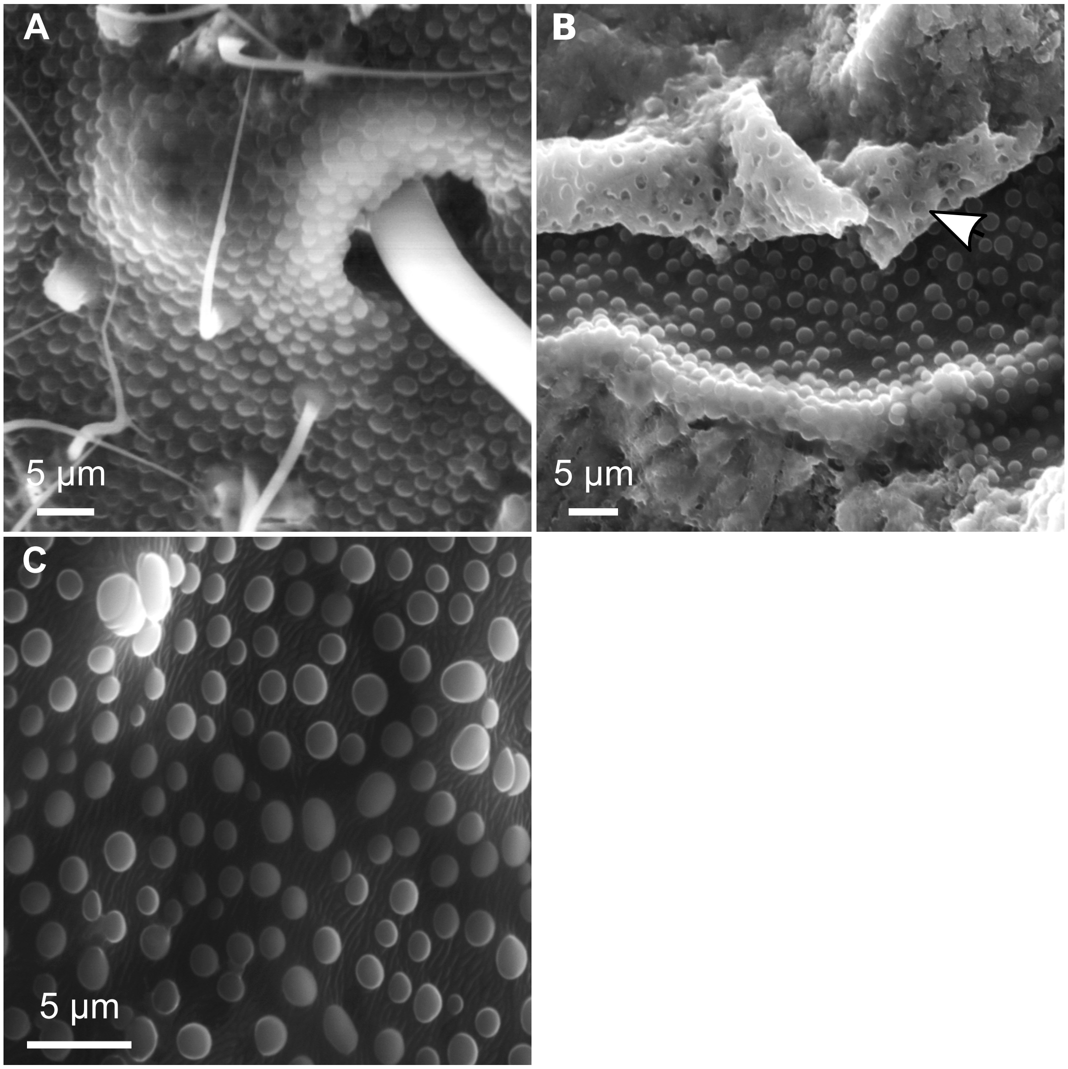


**Supplementary Fig. 7. Presence of a protein layer on the outermost exoskeleton of old worker of *Ac. echinatior*. (A)** Environmental scanning electron microscope (eSEM) image of original cuticle of *Ac. echinatior* showing papillae structures densely covering the epicuticle. **(B)** eSEM image of cuticle treated with 2.5 M KOH solution for 0.5 hour for dissolving surface proteins. The white arrow indicates the shedding protein layer with clear spores matching the papillae. **(C)** eSEM image of cuticle treated with 2.5 M KOH solution for 24 hours for complete dissolution of surface proteins uncovering the chitin skeleton.

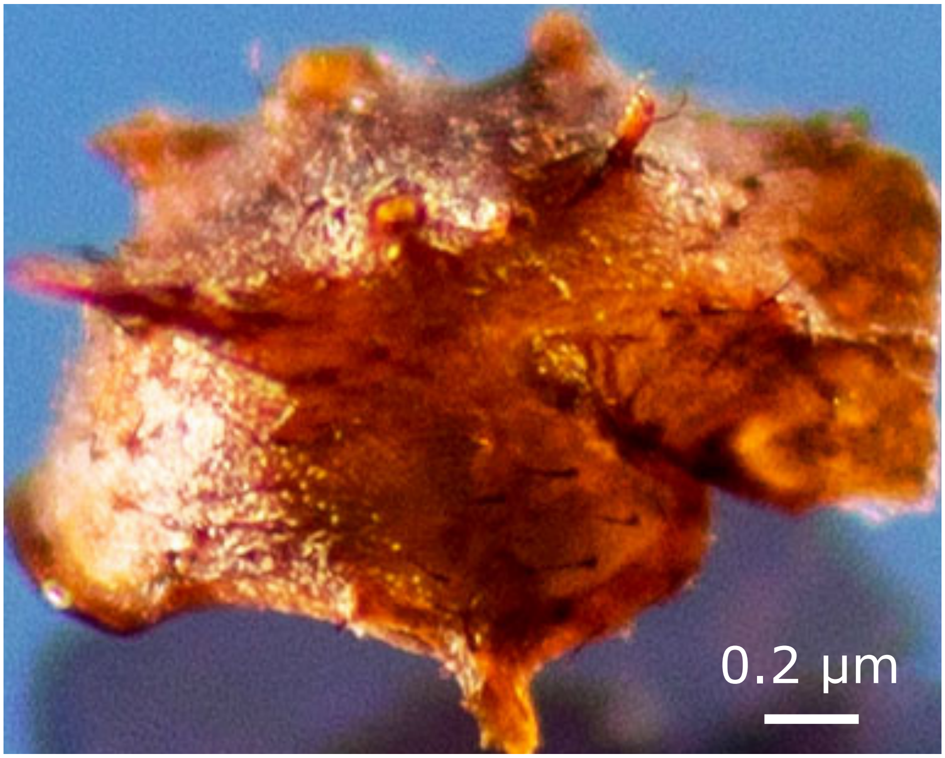


**Supplementary Fig. 8. The ant exoskeleton used for synthetic experiments.** The thorax part of ant exoskeleton used for synthetic biomineralization experiments after inside tissues have been carefully removed and the exoskeleton washed twice with distilled deionized water.

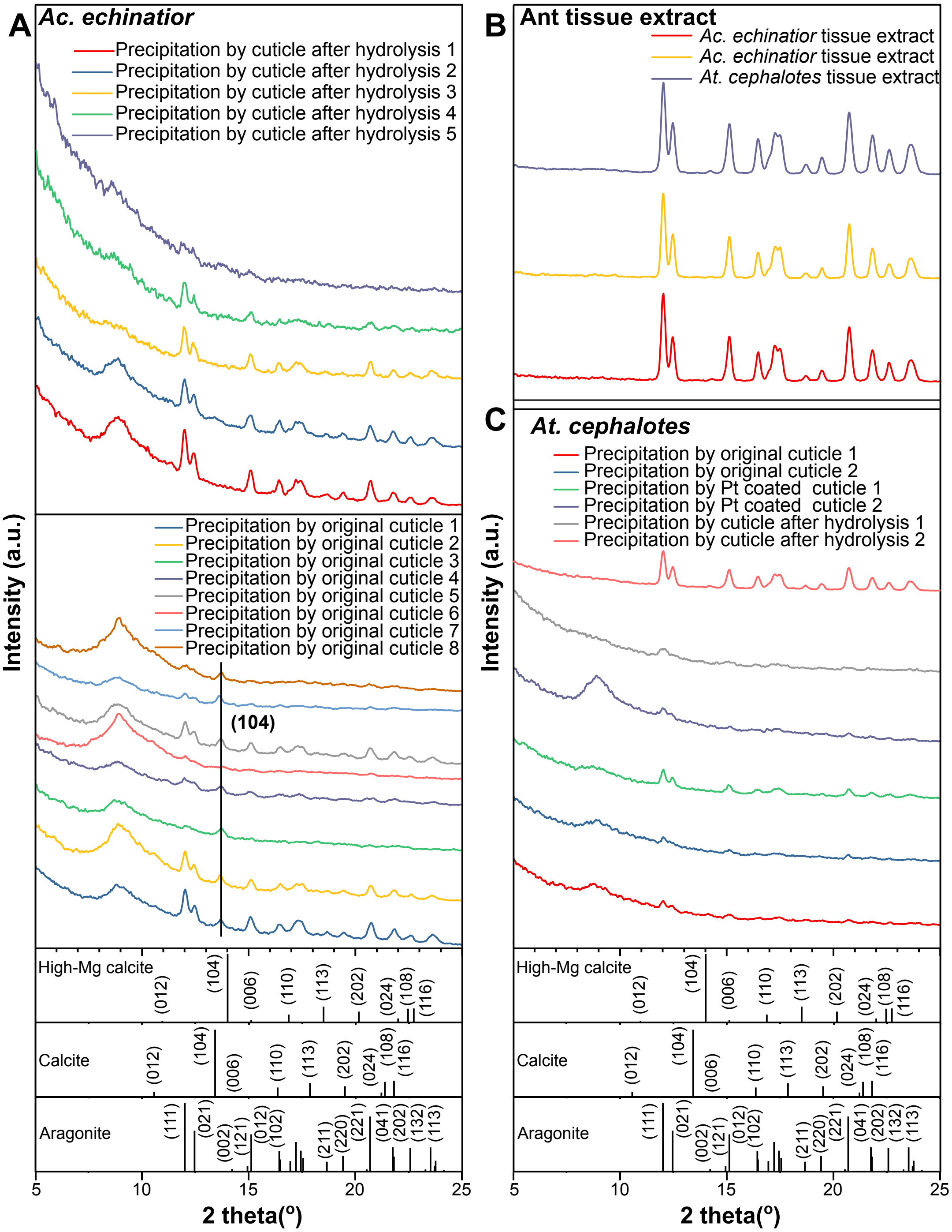


**Supplementary Fig. 9. X-ray diffraction analyses of *In vitro* synthetic experiments. (A)** XRD patterns of the synthetic carbonate induced by original *Ac. echinatior* cuticle, and for *Ac. echinatior* cuticle after KOH hydrolysis of protein layer. **(B)** XRD patterns of the synthetic carbonate induced by ant tissue extract. **(C)** XRD pattern of the synthetic carbonate induced by original cuticle, cuticle after KOH hydrolysis of protein and platinum coated cuticle of *At. cephalotes*.

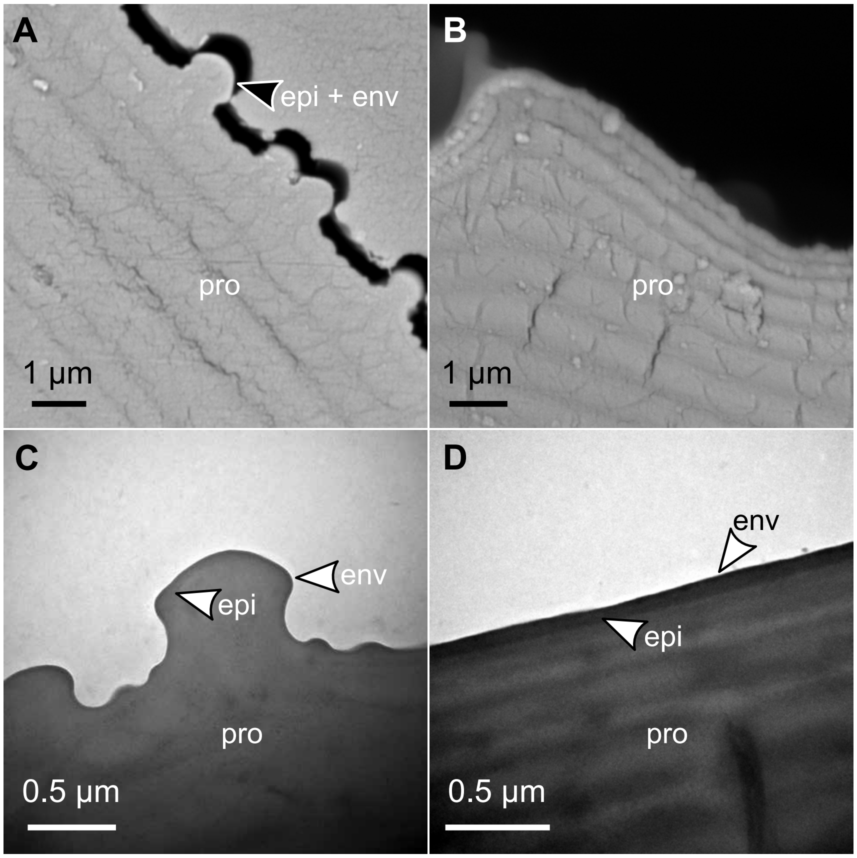


**Supplementary Fig. 10. Distinctly different cuticular structures present in *Ac. echinatior* compared to *At. cephalotes*.** SEM images of cross-sectioned cuticle showing **(A)** the characteristic lamellar procuticle (pro) covered by the epicuticle (epi) and a thin envelope (env) as the topmost layer in *Ac. echinatior,* whereas **(B)** the well-ordered lamellar structure of the exocuticle is significantly thinner in the *At. cephalotes* and the epicuticle and envelope layers cannot be distinguished. TEM images of cross-sectioned cuticle reveals **(C)** the presence of characteristic layers: the inner procuticle from the basis of the exoskeleton **(D)** with much darker epicuticle and envelope layers.

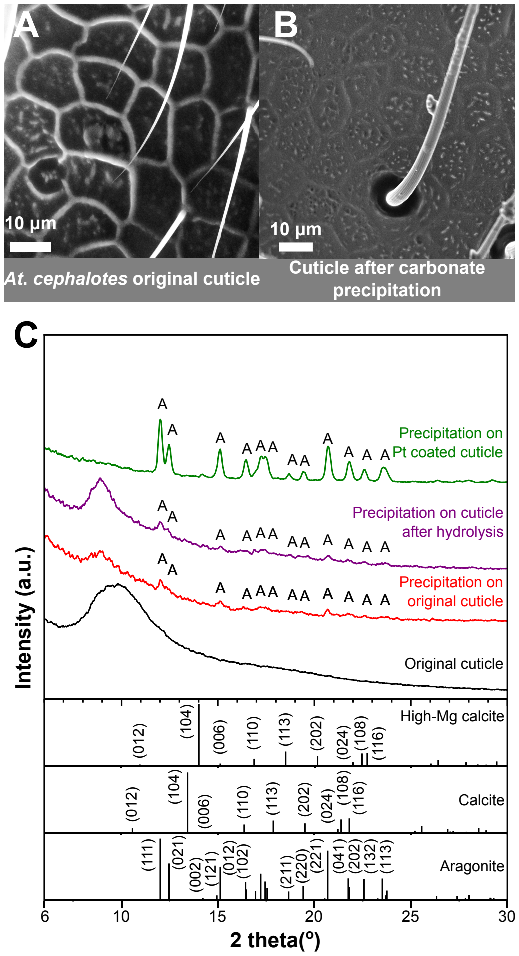


**Supplementary Fig. 11. SEM and X-ray diffraction analyses of *In vitro* synthetic experiments of *At. cephalotes*.** (A) SEM of original cuticle of *At. cephalotes*. (B) SEM of cuticle after carbonate precipitation experiment. (C)Typical XRD patterns of the synthetic carbonate induced by original cuticle, cuticle after KOH hydrolysis of protein and platinum coated cuticle of *At. cephalotes*, respectively. A: aragonite.

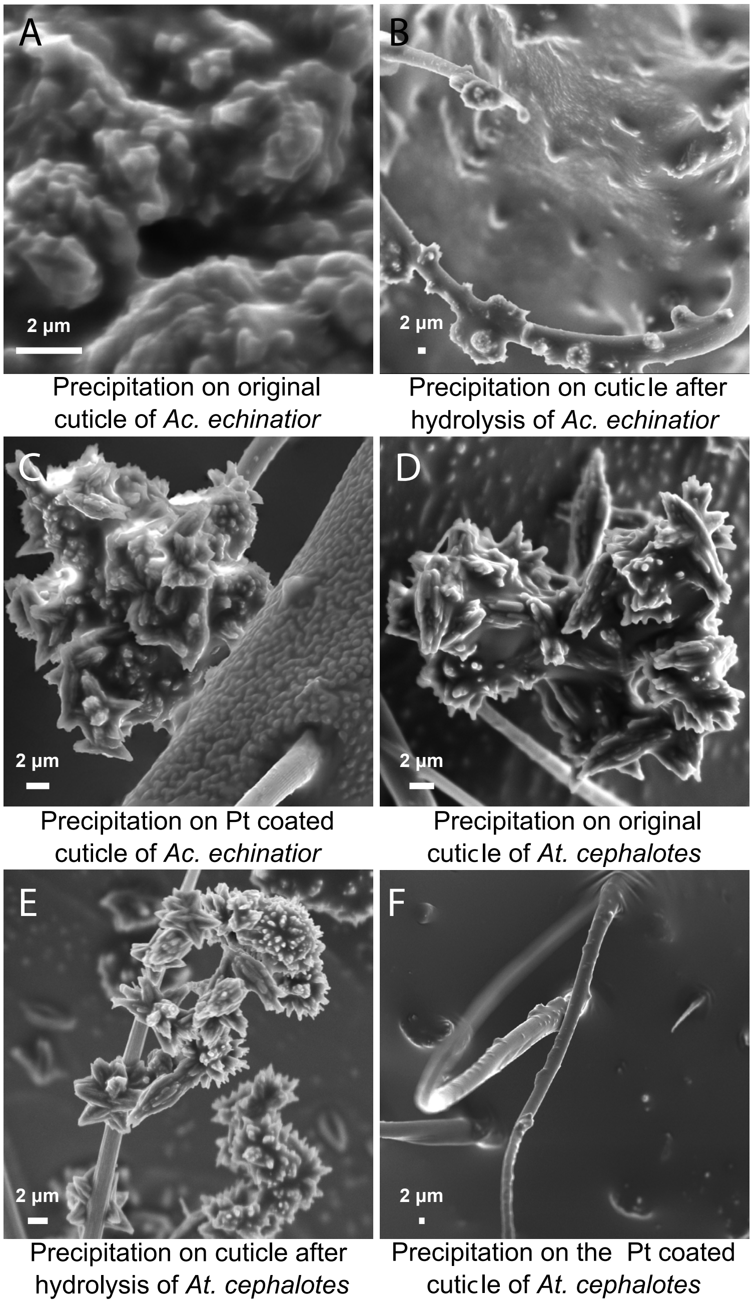


**Supplementary Fig. 12. Morphological and structural characterization of synthetic carbonates. (A, B, C, D, E,** and **F)** SEM micrographs obtained from the synthetic carbonate induced by original cuticle, cuticle after KOH hydrolysis of protein and platinum coated cuticle of *Ac. echinatior* and *At. cephalotes*. Consistent with the XRD pattern, high-Mg calcite only found in (A), while the rest show typical needle crystals of aragonite.

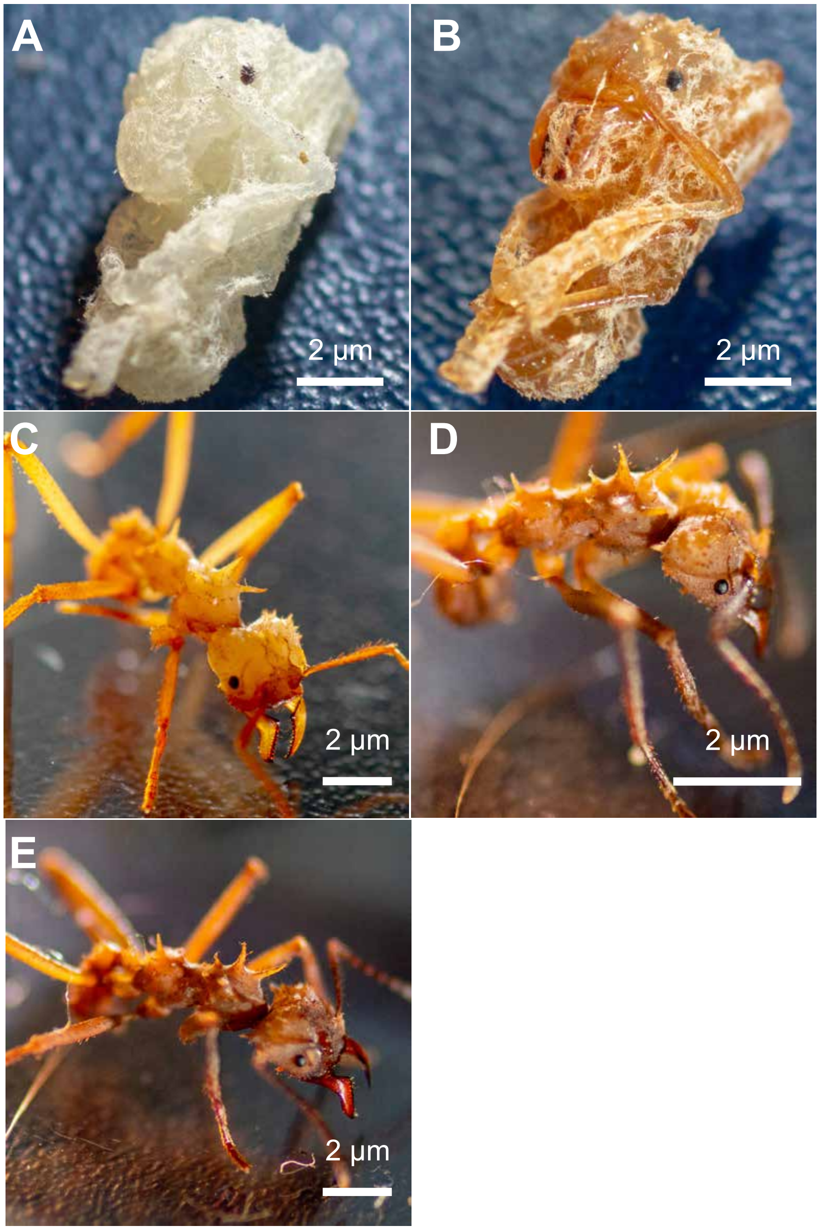


**Supplementary Fig. 13. Different cuticular coloration across development stages of *Ac. echinatior*.** New pupa **(A)**, pupa **(B)**, newly eclosed callow worker **(C)**, young worker **(D)**, and old worker **(E)**.

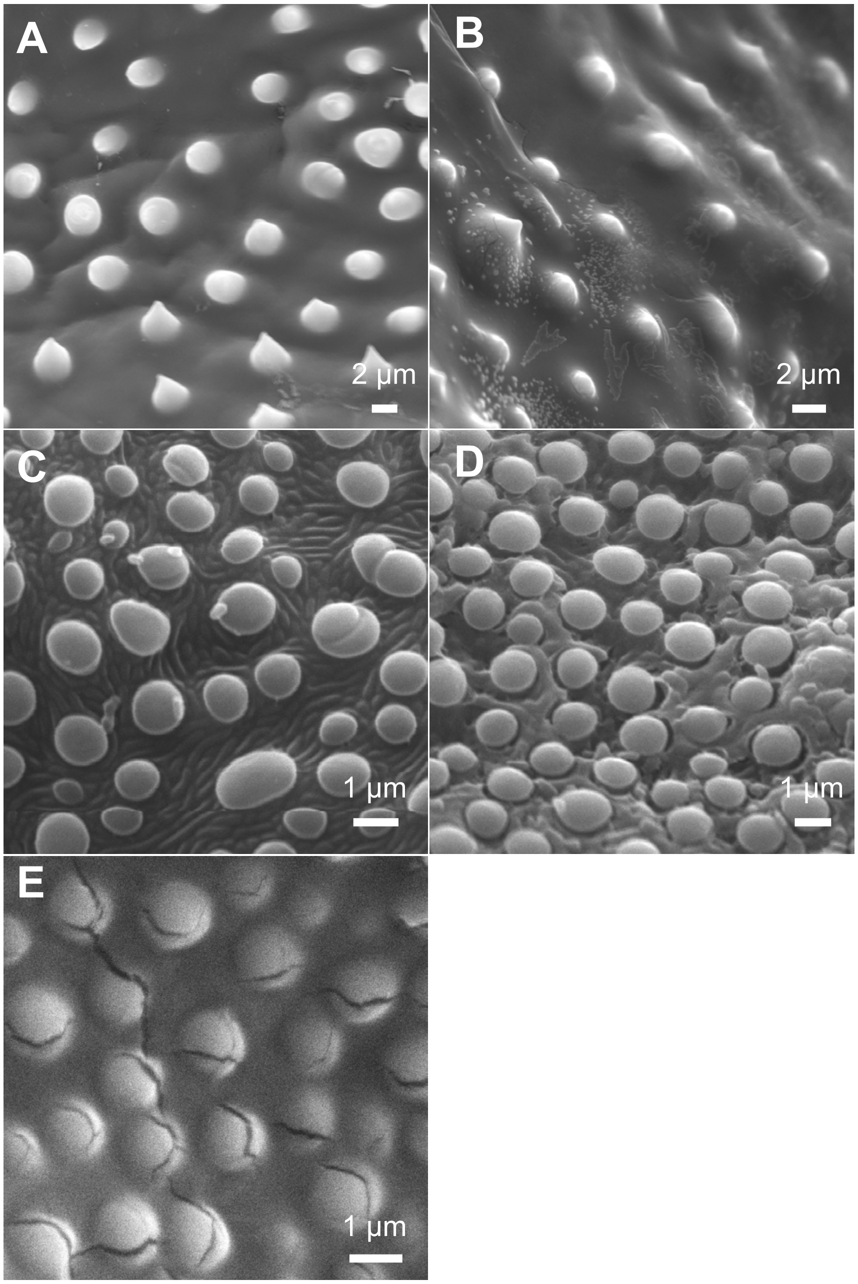


**Supplementary Fig. 14. Epicuticular difference across development stages of *Ac. echinatior*.** SEM images illustrating epicuticular papillae microstructures of new pupa **(A)**, pupa **(B)**, newly eclosed callow worker **(C)**, young worker **(D)**, and old worker **(E)**. Note that the increasing protein layer surrounding the epicuticular papillae represented along with the development stages, which is dissoluble by KOH solution as show above in Fig. S3.

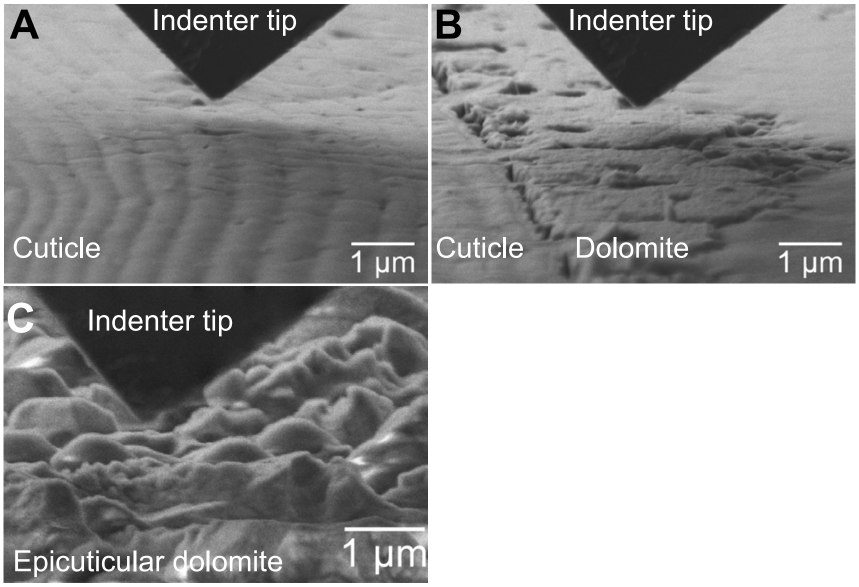


**Supplementary Fig. 15. SEM images illustrating *in-situ* nanoindentation measurements of ant cuticle hardness.** Nanoindentation on polished cross-sections of ant cuticle **(A)** and ant mineral **(B)** from the side**,** and unpolished epicuticular mineral-cuticle composite **(C)** from top-down. For the polished cross-sectional samples, the results represent hardness of the cuticle alone and the mineral alone, respectively, whereas the hardness measured on the unpolished composite sample from outside-in represents the combination of the two components. This is important since only by probing the cross-section from the side can we guarantee the "mineral alone" data actually represents the mineral itself, and not the mineral-cuticle composite. The irregularly shaped epicuticular mineral crystals on the outer surface can be clearly observed in **(C)**.

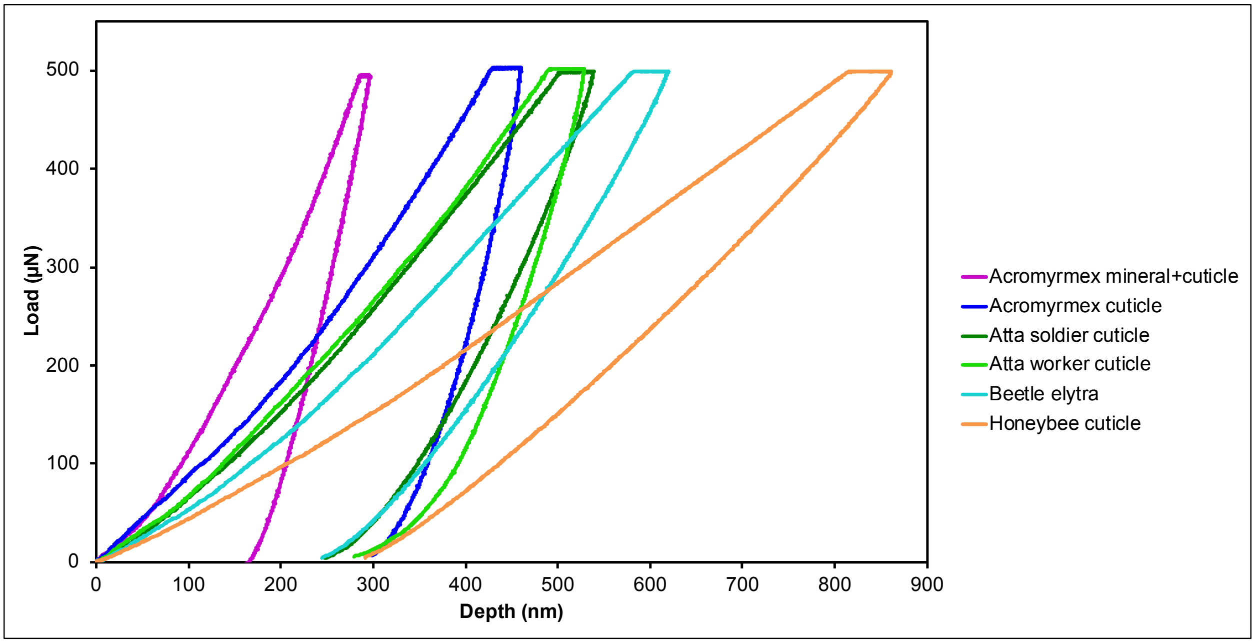


**Supplementary Fig. 16. Representative load-depth curves of samples measured with *in-situ* nanoindentation under a maximum load of 500 µN.** These curves illustrate different nano-mechanical responses from *Ac. echinatior* mineral + cuticle, *Ac. echinatior* cuticle, *At. cephalotes* soldier cuticle, *At. cephalotes* worker cuticle, beetle (*X. colonus*) elytra cuticle and honeybee (*A. mellifera*) cuticle.

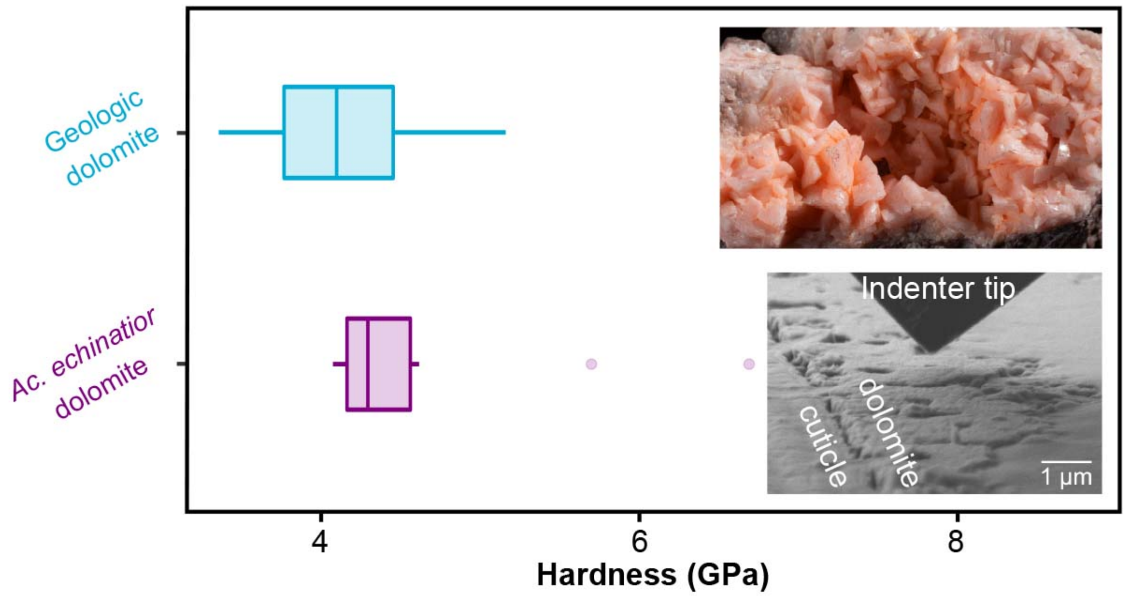


**Supplementary Fig. 17. Hardness of geologic dolomite and *Ac. echinatior* dolomite.** The dolomite formed on *Ac. echinatior* ants has an indentation hardness value of 4.59±0.73 GPa, similar to their geologic counterpart (4.08±0.49 GPa). Both were probed in *in-situ* nanoindentation on a polished flat sample.

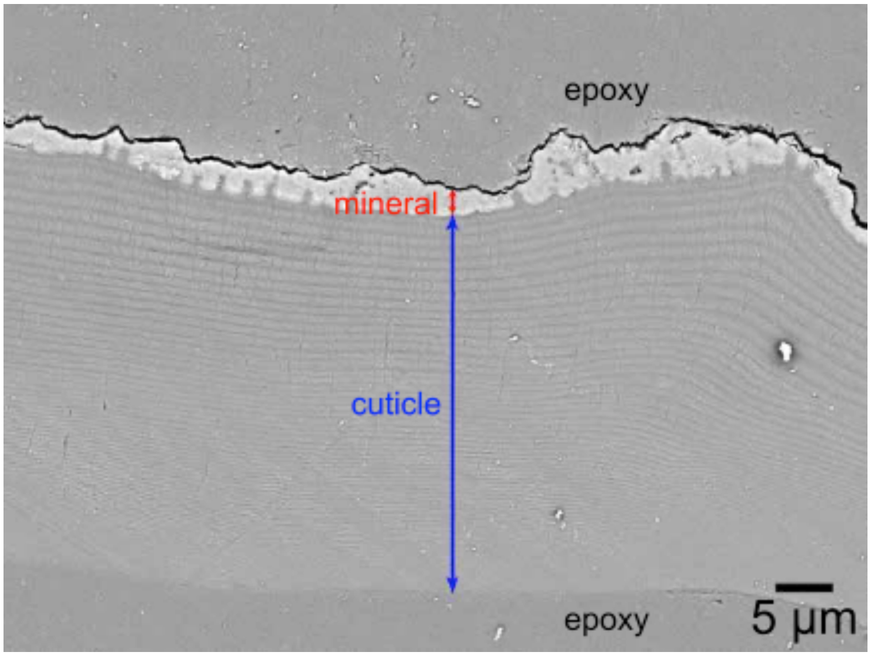


**Supplementary Fig. 18. Cross-sectional SEM micrograph of an adult *Ac. echinatior* cuticle.** Polished cross-section of the cuticle reveals the relative thickness of the mineral layer to the entire cuticle. Measurements of the thicknesses were taken on the exoskeletons across 5 areas from 3 different ants. The average thickness is 2.3 µm for the mineral layer and 33.5 µm for the cuticle, resulting in a thickness ratio of approximately 0.07.

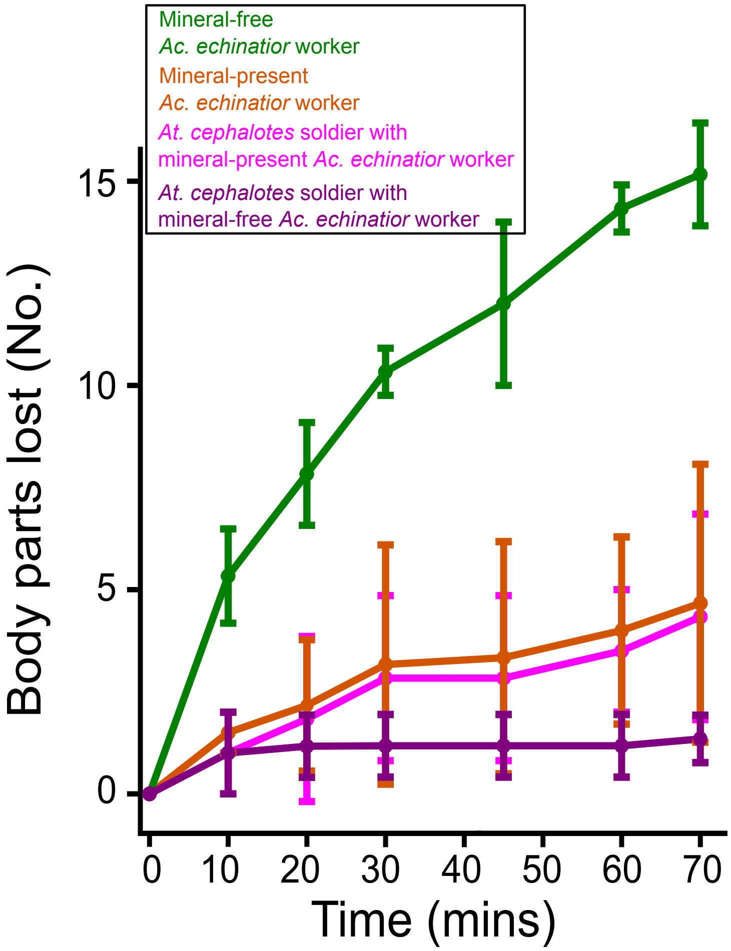


**Supplementary Fig. 19. Number of body parts lost during the ant battle tests between *Ac. echinatior* major worker and *At. cephalotes* soldiers.**

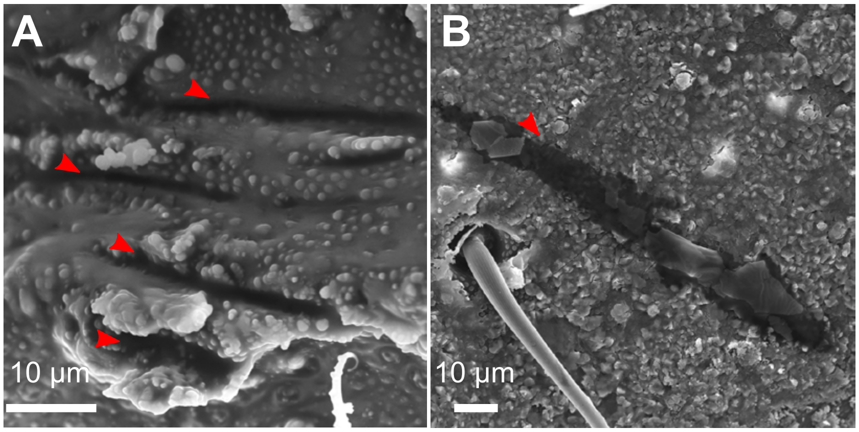


**Supplementary Fig. 20. SEM images of wounds inflicted on *Ac. echinatior* major worker by *At. cephalotes* soldiers.** Compare to the mineral-free ants **(A)**, the mineral ant showed significantly less damage to their exoskeleton **(B)**. The red arrows indicate bite marks by *At. cephalotes* soldiers.

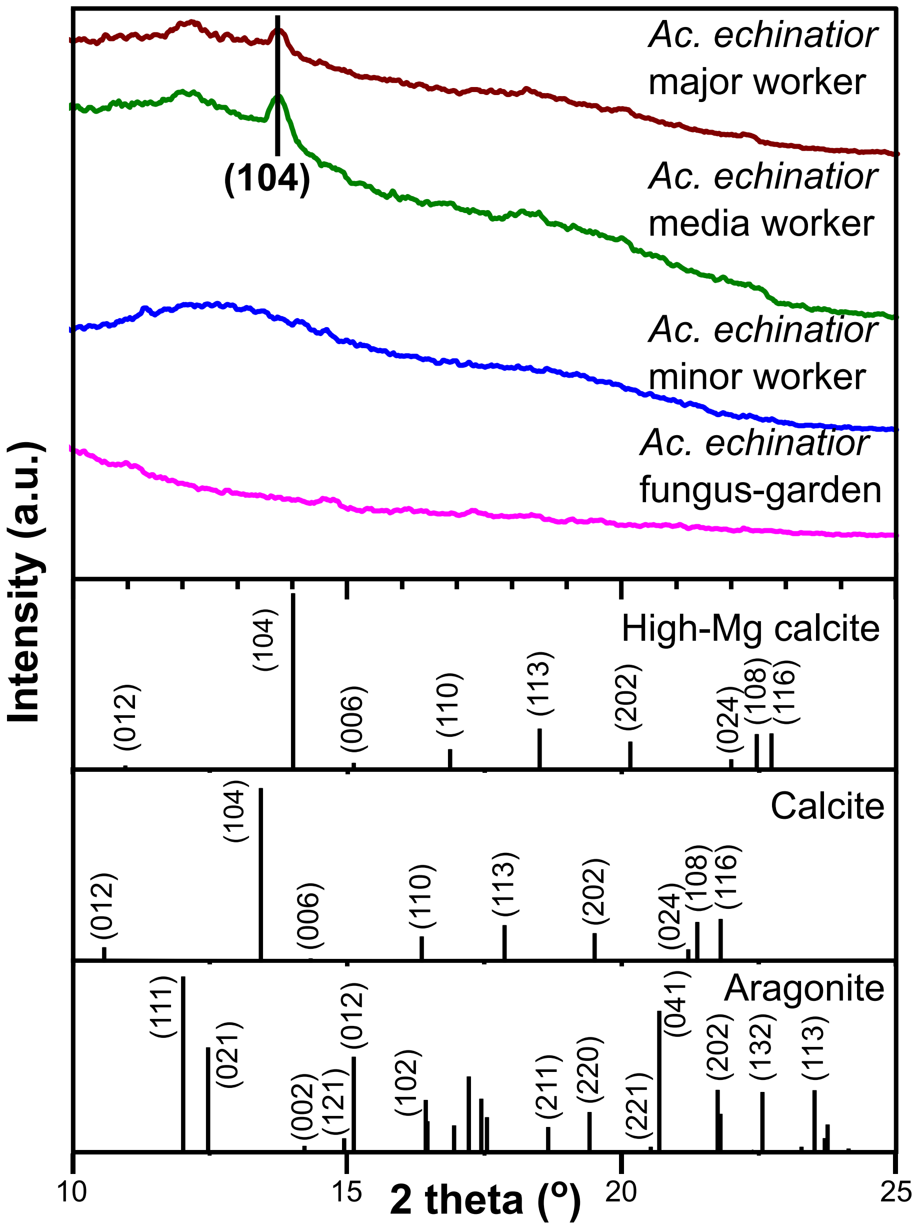


**Supplementary Fig. 21. Presence and absence of mineral armor in different worker castes of *Ac echinatior* and in fungus garden.** Only media and major worker have mineral armor as determined by XRD analyses.

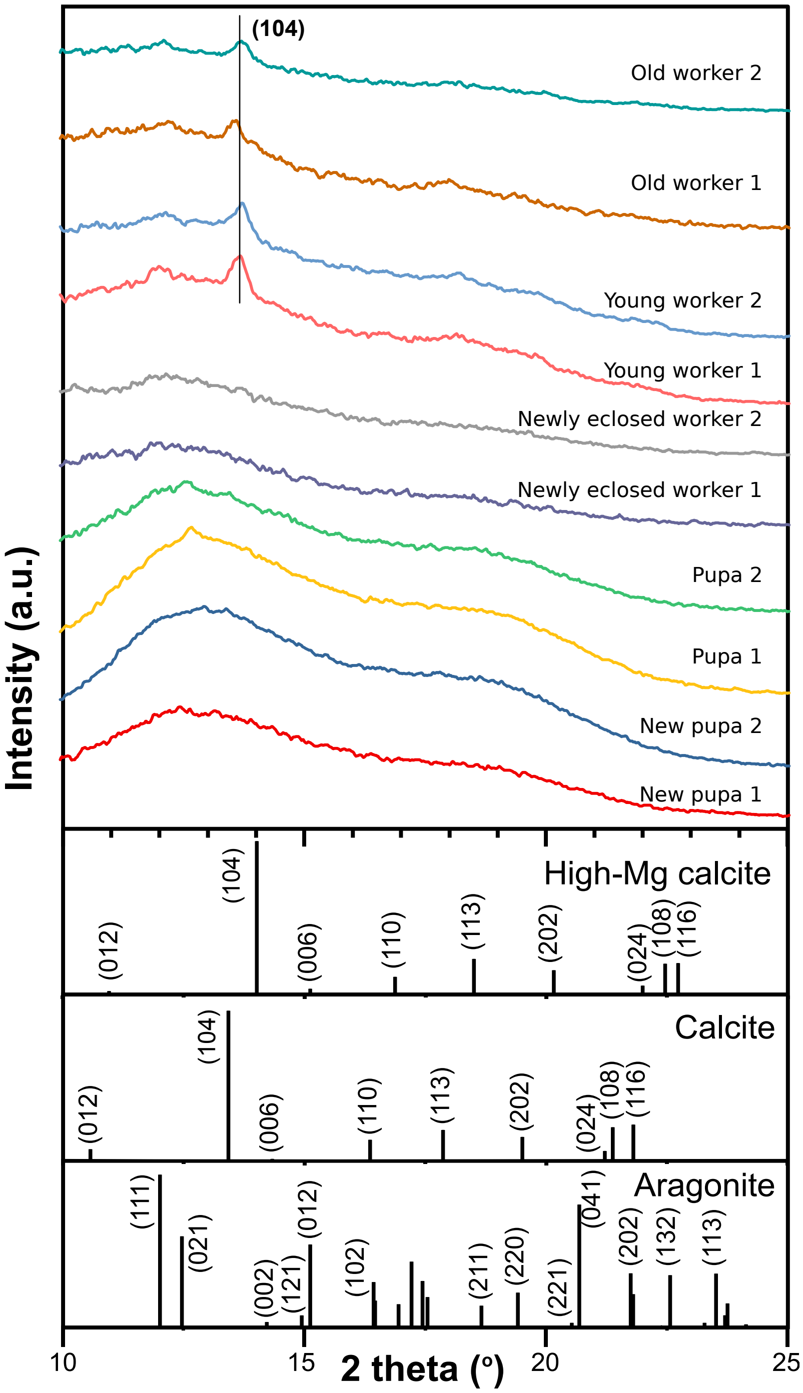


**Supplementary Fig. 22. Presence and absence of mineral armor in *Ac. echinatior* major workers and pupae of different ages.** Results show that only mature workers (young and old workers) have mineral armor as determined through XRD analyses.

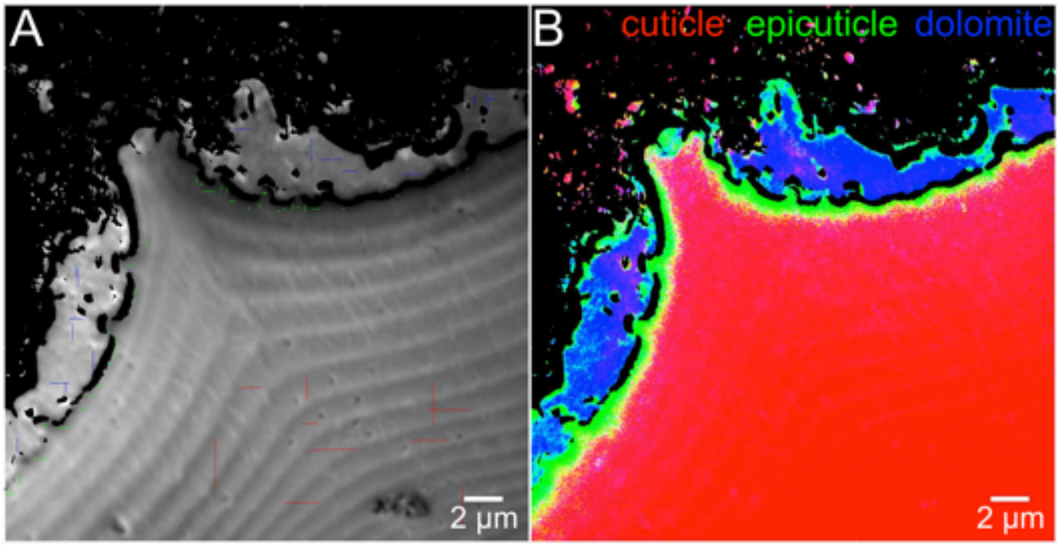


**Supplementary Fig. 23. PEEM images. (A)** Average of 181 PEEM images acquired across the C K-edge (280-320 eV). Colored lines indicate the pixels from which 500 or more single-pixel spectra were extracted for each component: red lines in the cuticle region, green lines in the epicuticle region, and blue lines in the mineral region. The 500 single-pixel spectra were extracted from each region, aligned, and then averaged to provide the 3 component spectra displayed in Fig. 2b. **(B)** A fully quantitative RGB component map of the same region as in Fig. 2c, with the blue channel not enhanced 5×, thus the intra-cuticle and intra-epicuticle mineral particles appear a faint magenta. In the enhanced map in Fig. 2c they are blue or purple.

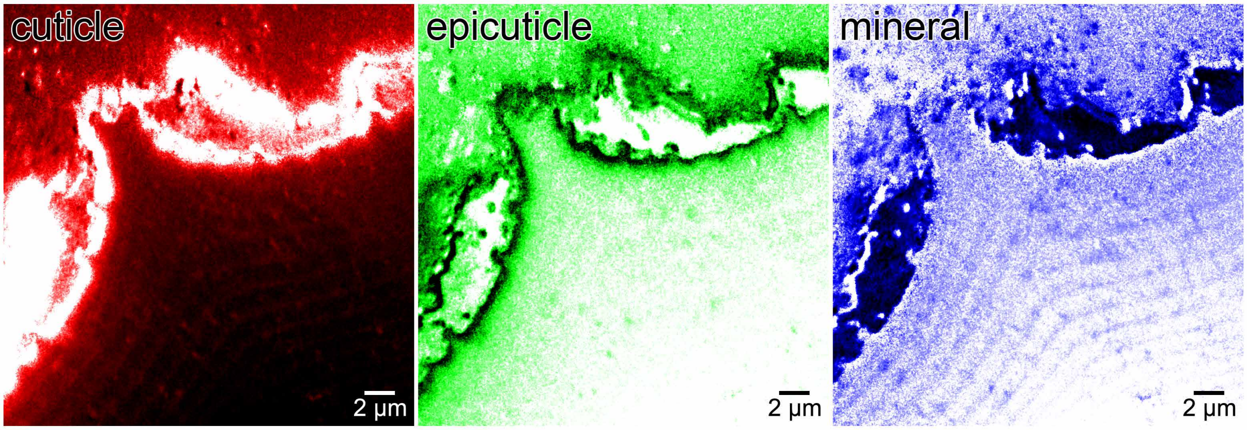


**Supplementary Fig. 24. Individual component maps from PEEM analysis.** Individual distribution maps of the three components characterized from PEEM: cuticle, epicuticle, mineral. Color-coded as in Fig. 1e. Background is inversed in the mineral component map, in order to better visualize the carbonate gradients in the cuticle, with darker colors indicating higher concentration (i.e., white represents 0% and black represents 100% of each component).

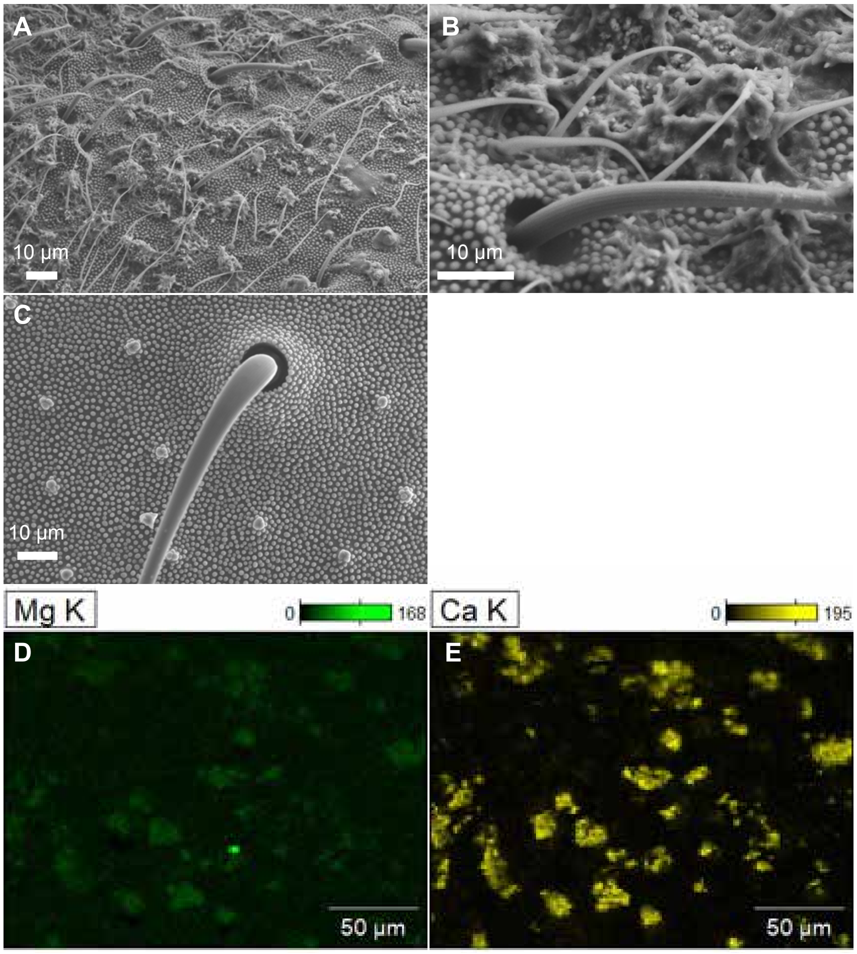


**Supplementary Fig. 25. Magnesium- and calcium-rich secretion on the cuticle of young worker ants of *Ac. echinatior*.** SEM image of mineral -free *Ac. echinatior* ant revealed abundant secretions scatter on the epicuticle **(A** and **B)**. After removing these secretions using mouthwash solution for 1 hour (Li et al., 2018 *PNAS*)[16], the presence of the specialized tubercle structures and papillae are revealed over the cuticle surface **(C).** Taken together, with our previous studies showing that each tubercle is connected to an internal glandular cell via a channel through the exoskeleton (Currie et al. 2006 *Science*)[18], our work suggests these secretions are ant gland-derived. EDS analyses show that the glandular secretions are enriched with magnesium and calcium ions **(D** and **E)**.

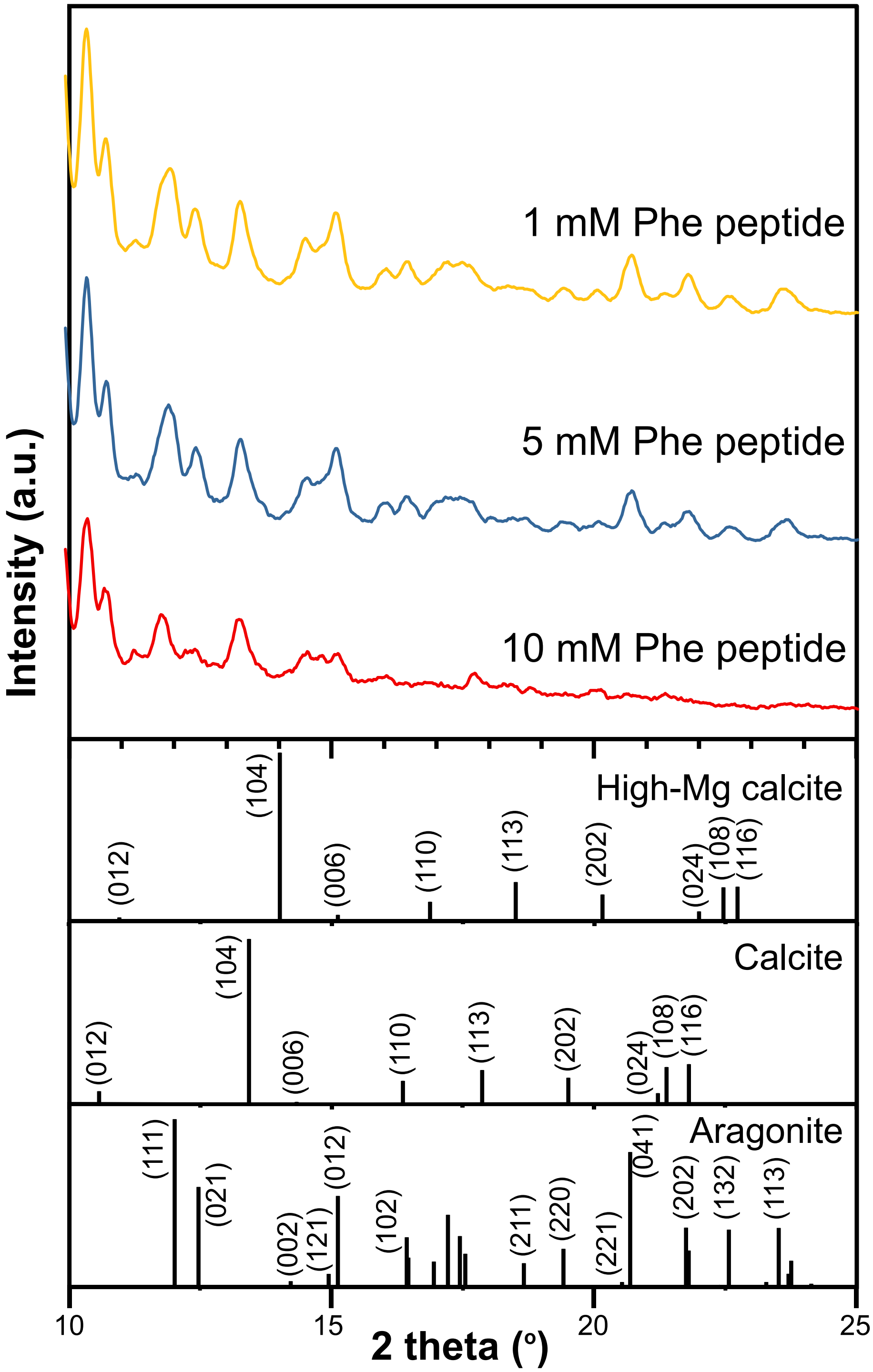


**Supplementary Fig. 26. Synthetic biomineralization experiments using phenylalanine-rich peptides with different concentrations.** Exclusively aragonite is found in the synthetic experiments using (H-Phe-Phe-Phe-OH) peptide as XRD analysis.

**Supplementary Video 1.**

Representative real-time microscopic video of *in-situ* nanoindentation measurement on *Ac. echinatior* cuticle.

Supplementary Video 2.

The time lapse of aggressive experiments between *At. cephalotes* soldier and three *Ac*. *echinatior* worker ants with mineral-present or absent.

**Supplementary Video 3.**

Details of aggression experiments between *At. cephalotes* soldier and mineral-free *Ac*. *echinatior* worker ants. Note that *At. cephalotes* soldier cut throughout thorax of mineral-free *Ac*. *echinatior* worker ants.
